# Spatiotemporal dispersion of DENV1 genotype V in western Colombia

**DOI:** 10.1101/2024.07.24.605015

**Authors:** Diana Rojas-Gallardo, Tyshawn Ferrell, Paula Escobar, Diego Lopez, Beatriz Giraldo, Juliana Restrepo-Chica, Erika Jimenez-Posada, Marlen Martinez-Gutierrez, Julian Ruiz-Sáenz, Autum Key, Nima Shariatzadeh, Dara Khosravi, Megan A. Martinez, Andrei Bombin, Jesse J. Waggoner, Jorge E. Osorio, Christopher J Neufeldt, Matthew H Collins, Jaime A. Cardona-Ospina, Anne Piantadosi

**Affiliations:** Population Biology, Ecology and Evolution Graduate Program, Emory University, Atlanta, GA, USA; Grupo de Investigación Biomedicina, Facultad de Medicina, Institución Universitaria Visión de las Américas, Pereira, Colombia; Biochemistry, Cell & Developmental Biology Graduate Program, Emory University, Atlanta, GA, USA; Grupo de investigación en Inmunología Molecular, Facultad de Ciencias de la salud, Maestría en Ciencias Biomédicas, Universidad del Quindío, Colombia; Facultad de Ciencias de la Salud, Unidad Central del Valle del Cauca, Tuluá, Colombia; Instituto para la investigación en Ciencias Biomédicas – Sci-help, Colombia; Grupo de Investigación en Ciencias Animales, Universidad Cooperativa de Colombia, Bucaramanga, Colombia; Department of Pathology and Laboratory Medicine, Emory University School of Medicine, Atlanta, Georgia, USA; Microbiology and Molecular Genetics Graduate Program, Emory University School of Medicine, Atlanta, Georgia, USA; Division of Infectious Diseases, Department of Medicine, Emory University School of Medicine, Atlanta, Georgia, USA; Department of Pathobiological Sciences, School of Veterinary Medicine, University of Wisconsin, Madison, WI, USA; Department of Microbiology and Immunology, Emory University School of Medicine, Atlanta, Georgia, USA; Division of Infectious Diseases and Vaccinology, School of Public Health, University of California, USA

**Keywords:** DENV-1V, Spatiotemporal dispersion, Phylogenetic analysis, Genetic diversity, Colombia

## Abstract

Dengue virus (DENV) is a significant public health concern in Colombia, with increased transmission of DENV type 1 (DENV-1) in the departments of Risaralda and Valle del Cauca in the Central-West region of the country following a large outbreak in 2019. However, little is known about the source, genetic diversity, and evolution of circulating viruses. We obtained plasma samples from individuals with acute DENV infection and analyzed DENV-1 genetic diversity, phylodynamics, and phylogeography. We found that most viruses belonged to DENV-1 genotype V, and phylogenetic analysis revealed three distinct clades, each of which was most closely related to viruses from neighboring departments of Colombia sampled over the last 5-10 years. Thus, the 2019 outbreak and subsequent DENV-1 circulation was not due to the introduction of a new lineage but rather reflected local DENV-1V dispersion and evolution. We identified amino acid positions under positive selection in structural proteins and NS1, which may have a role in immune evasion and pathogenesis. Overall, our analysis of DENV1 genotype V diversity, evolution and spread within Colombia highlights the important role of genomic surveillance in understanding virus dynamics during endemic circulation and outbreaks.

## Introduction

Dengue virus (DENV) is estimated to cause 390 million infections globally each year, primarily in tropical and subtropical regions [1, 2]. Infection can cause mild symptoms, such as fever, myalgia, and headache, or severe syndromes, including dengue hemorrhagic fever (DHF) and dengue shock syndrome (DSS). DENV belongs to the *Flaviviridae* family and is transmitted by *Aedes aegypti* (primary) and *Aedes albopictus* (secondary) mosquito vectors [3, 4]. The global distribution of DENV is determined primarily by the geographical range of these mosquito species, but other factors, such as the genetic diversity of the virus, the host population’s immunity, and socio-ecological (i.e. urbanization) conditions, also influence transmission [5, 6]. The Pan American Health Organization (PAHO)’s most recent epidemiological update indicates that the total number of cases in the Americas has increased over 20-fold since the 1980s, especially within South America [7, 8]. Additionally, DENV-related mortality has increased over 2-fold in the Americas in the past five years [9, 10]. However, the extent of DENV ‘s impact in the Americas is likely not fully appreciated due to limited surveillance.

In addition to diagnostic and epidemiological surveillance, genomic surveillance can be crucial to understanding patterns of dispersion and evolution during both outbreaks and endemic circulation. For example, genomic surveillance can detect new introductions, reveal transmission hotspots, and identify emerging variants with increased transmissibility. This information can, in turn, inform targeted strategies for DENV control. Here, we used genomic epidemiology to investigate DENV transmission and evolution in western Colombia, which has experienced an increasing burden of disease since 2019.

DENV is a positive-sense RNA virus that has a genome length of 10.7kb and encodes three structural proteins (capsid, pre-membrane, and envelope) and seven nonstructural proteins (NS1, NS2A, NS2B, NS3, NS4A, and NS5) [11, 12]. DENV comprises four antigenically distinct serotypes (DENV 1-4) whose envelope proteins have an amino acid similarity of only about 70% [13]. Within each (sero)type, there are multiple distinct genotypes; viruses share ≤94% genetic similarity between genotypes and ≥95% genetic similarity within genotypes [14-18].

DENV is the most prevalent arbovirus in Colombia and is thought to have been introduced from the Lesser Antilles in the early 1970s [19, 20]. DENV-2 was introduced first, followed by DENV-3, DENV-1, and DENV-4[21]. The four DENV types have been co-circulating since the early 1990s, though DENV-1 is currently predominant [22]. DENV-1, which is generally associated with milder symptoms than other types, comprises five genotypes distributed across distinct geographic regions [23, 24]. Genotype V has been the most predominant in the Americas over the past several decades, after replacing genotype III [25]. In addition to endemic circulation, each DENV type has also been reported in region-dependent outbreaks across the country [26]. Colombia has six regions: Andean (Central), Amazonia, Pacific Coast, Orinoco, Caribbean Coast and Insular Region. The Central-West region has had five major DENV outbreaks: in 1989, 2002, 2010, 2013, and 2019 [21].

In 2019, starting from the eighth epidemiological week, Colombia experienced a new epidemic phase of dengue in which the reported cases were above the upper limit compared to its historical behavior (2011-2018) and the accumulated incidence was 465.9 cases per 100,000 residents at the end of the year [27]. Since then, the number of DENV cases in Colombia has nearly doubled, and severe dengue increased over 2-fold in 2023 [9].

We hypothesized that viral factors may have contributed to this increased disease burden, for example the introduction of new lineages and/or mutations that conferred immune evasion. Prior studies have highlighted a role for clade replacement in driving DENV outbreaks in other locations [28-30], and Salvo *et al*. described both serotype shift (from DENV1 to DENV3) and within-DENV1 lineage replacement in the Colombian department of Antioquia from 2014-2016 [31, 32]. However, DENV transmission dynamics and evolution in the Central-West region of Colombia remain unknown, despite the high incidence of DENV in this region.

We generated new DENV surveillance and sequence data from 2019 to 2022 to investigate the origin, genetic diversity, and spatiotemporal dispersion of DENV lineages into and across departments in Colombia. We particularly focused on two nearby cities in the Central-West region with different disease burdens, Cali located in Valle del Cauca (303 cases per 100,000 residents in 2021) and La Virginia located in Risaralda (18 cases per 100,000 residents in 2021) (**Figure S1**). We focused on DENV-1 genotype V (DENV-1V), which was the most common virus identified in our contemporary sequences, and which had previously been very little studied, with only about 30 full-length sequences from Colombia published in GenBank since 1988. Prior to our study, only 1 DENV genome sequence had been generated from the departments of Risaralda and Valle del Cauca, underscoring the need for increased genomic surveillance in this region. Our new sequence data allowed us to investigate patterns of dispersion and evolution within the Central-West region and place this in context of DENV dynamics within the country and region.

## Methods

### Ethics Statement

The acute febrile illness study in La Virginia, Risaralda, is approved by the ethics committee of the Universidad de Cooperativa de Colombia. Samples collected in Cali, Valle del Cauca, were remnant clinical specimens, and this study has the approval of the ethics committee of the Universidad Unidad Central del Valle del Cauca UCEVA. Work performed at Emory University was considered secondary use of deidentified samples, and no IRB review was required.

### Study Locations, including DENV burden

Colombia is administratively divided into 32 departments (First administrative level). In accordance with the guidelines of the national public health surveillance system, all probable and confirmed cases of dengue require mandatory reporting to the National Public Health Surveillance System (SIGIVILA). This process includes data consolidation and report from the local institution to the departmental institution, and then analysis and publication by the national health institute (INS).

The municipality of La Virginia belongs to the department of Risaralda in the central west part of Colombia and has a total population for 2023 of 28,488 inhabitants. La Virginia, referred to by its department Risaralda hereafter, is located approximately 30 km from the state capital, Pereira. It is 200 km North of Cali, the capital of the department of Valle del Cauca, and 210 km south of Medellín, the capital of the department of Antioquia. Risaralda has endemic DENV circulation and reported 153 cases in 2019, 125 cases in 2020, 26 cases in 2021, and 22 cases in 2022 to SIVIGILA (SIVIGILA consulted in August 2023).

The city of Cali, located in southwestern Colombia, is the third largest city in the country and has a total population for 2023 of 2,280,522 inhabitants. Cali, referred to by its department Valle del Cauca hereafter, has endemic circulation of dengue virus with seasonal outbreaks and reported a total of 3,889 cases in 2019 (3.1% of the national total), 13,232 cases in 2020 (16.9% of the national total), 5,726 cases in 2021 (11% of the national total) and 2,840 cases in 2022 (4.1% of the national total) (data from data the epidemiological bulletins, Instituto Nacional de Salud INS).

### Sample collection

Serum samples were collected from patients in a prospective cohort study in Risaralda between July 2019 and April 2022. In this cohort, patients residing in the Central Western Metropolitan Area of Risaralda were recruited at the San Pedro y San Pablo Hospital if they presented with a febrile condition that was diagnosed as probable dengue. A trained nurse was responsible for informed consent and assent, verifying the inclusion and exclusion criteria. Clinical assessment and classification were performed by the attending physician, and data was obtained from clinical records and a structured questionnaire.

Serum samples from patients in Valle del Cauca were obtained from August 2021 to December 2022 at a medical center located north of the city. These were residual clinical samples from patients diagnosed with dengue by clinical laboratory testing.

### DENV-1 detection and typing

Samples from both Risaralda and Valle del Cauca were transported to the molecular biology laboratory at the Universidad Vision de las Américas, and a rapid test was used to detect the NS1 antigen and IgG and IgM envelope-specific antibodies (Bioline™ Dengue Duo). RNA was extracted from samples positive for NS1 and IgM using the PureLink™ RNA Mini Kit (Invitrogen). To confirm dengue infection and detect infections with other arboviruses, such as Zika and chikungunya, a multiplex RT-qPCR assay was performed using a previously described assay [33]. PCR results were confirmed for all samples by testing a duplicate aliquot using the same assay at Emory University. In RT-qPCR positive samples, the dengue serotype was determined by a multiplex real-time RT-qPCR method as previously described [34].

### Full DENV genome sequencing

We used two approaches to generate full-length DENV genome sequences: metagenomic and multiplex amplicon sequencing. For both approaches, we first performed heat-labile dsDNase treatment (ArcticZymes, Tromso, Norway) and first-strand cDNA synthesis with SUPERSCRIPT IV RT and random primers (Fisher/Invitrogen, Hampton, USA).

### Metagenomic sequencing and analysis

For metagenomic sequencing, second-strand cDNA synthesis was performed using NEB reagents, and double-stranded cDNA underwent Nextera XT library construction and Illumina sequencing as previously described [35]. As a positive control, External RNA Controls Consortium (ERCC) spike-ins (Illumina) were added prior to cDNA synthesis. The ERCC spike-ins also served as a unique control for each sample to measure cross-contamination. At least one negative control (water) was included with each library construction batch. After Illumina sequencing, adapter trimming was conducted in BaseSpace using bcl2fastq2 v2.20 to mask short adapter reads (n = 35) and trim to a minimum read length (n = 35). Reads were processed using viral-NGS v1.25.0 [36] in the following manner: they were converted to unmapped bam files, quality control checked for the correct ERCC spike-in, and taxonomically classified using KrakenUniq. Reads underwent reference-based assembly, initially using the DENV1 RefSeq sequence (NC_001477). The resulting partial genomes were evaluated by BLAST to find the best-matching reference sequence, OM654348, which was used for final reference-based assembly, all using viral-NGS v1.25.0 [36].

### Multiplex amplicon sequencing primer design

We designed a new multiplex amplicon sequencing protocol to specifically sequence lower-concentration DENV1 genotype V viruses from this study based on the sequence data we had generated by metagenomic sequencing (**Figure S2**) [37]. Multiplex amplicon primers were designed using a multiple sequence alignment of five DENV1 Genotype V sequences we had generated by metagenomic sequencing, which had coverage of >99% and mean depth of >200x and represented different departments of Colombia. PrimalScheme v1.4.1 [38] was used to generate 34 primer pairs that amplified ∼400 nucleotide (nt) overlapping fragments spanning the entire genome. These primers were mapped to our sequences to check for mismatches and amplicon lengths. Any primers that had multiple binding sites or could lead to primer-dimer formation were modified. Primers with a Tm above 50.1°C (ranging from 55.9 -61°C) were selected to reduce self-dimer and hairpin secondary structures. The amplicon primers from PrimalScheme were concatenated with Nextera XT indexes to create fusion primers, which contained a forward iNEXT universal primer (5’ TCGTCGGCAGCGTCAGATGTGTATAAGAGACAG 3’) at the 5’ end of the forward primer and a reverse iNEXT universal primer (5’ GTCTCGTGGGCTCGGAGATGTGTATAAGAGACAG 3’) at the 5’ end of the reverse primer, as shown in (**Table S1**) [39]. The fusion primers were synthesized by Integrated DNA Technologies (Iowa, USA).

### Multiplex amplicon library construction and sequencing

To generate DENV1 amplicons from the first-strand cDNA template described above, the DENV-specific multiplex primers were separated into two pools (odd and even numbers from **Table S1**), and multiplex PCR was performed in two separate reactions per sample to reduce the overextension of neighboring amplicons. Each multiplex PCR reaction contained cDNA template (3 ul), Q5 Hot Start high-fidelity DNA polymerase (12.5 ul) (NEB), nuclease-free water (7.5 ul), and the odd or even primer pool (2 ul). The multiplex PCR cycle parameters were set as follows: 98°C for 30 seconds, 30 or 34 cycles at 95°C for 15 seconds, and 65°C for 5 minutes. The cycle number depended on the DENV C_T_ value: 30 cycles were used for samples with a C_T_ value ≤ 30, while 33 cycles were used for samples with a C_T_ value above 30. The amplicons underwent a thirteen-cycle Nextera indexing PCR using dual unique indexes. Resulting libraries were quantified using the KAPA Universal Complete Kit (Roche), pooled to equimolar concentration, and sequenced using a Miseq 600 cycle v3 kit (Illumina) with paired-end 300bp reads. The median number of reads per sample was 3.03M (range: 882K - 8.47M).

### Reference-based assembly

FASTQ files from multiplex amplicon sequencing were analyzed using ViralRecon v2.6.0 [40]. Reads were filtered based on the Q30 score, length (minimum 50 bp), and Qphred score (minimum 30) using FastQC v0.11.9. The reads underwent adapter and quality trimming before removing host reads using Fastp v0.23.2 and Kraken 2 v2.1.2. Next, the paired-end reads were mapped to the reference sequence (OM654348) using Bowtie 2 v2.4.4. Consensus sequences were then assembled in ViralRecon.

### Optimizing multiplex amplicon primer pooling

To optimize the concentration of each primer in our multiplex amplicon pools, we first sequenced 3 samples that had previously been sequenced using the metagenomic approach, measured the per-amplicon depth using mosdepth v0.3.3, and calculated a sequence read fraction (SRF) by dividing the amplicon’s depth by the median depth across the whole genome. Amplicons with an SRF value between 0.75-1.3 were deemed adequate. For amplicons with SRF values lower or higher than this range, we increased or decreased the primer concentrations, respectively, and repeated testing. The final primer concentrations are listed in **Table S1**.

### Alignment with reference sequences

All DENV-1V full genome and envelope gene sequences were downloaded from NCBI, and their genotypes were verified using Genome Detective v3.83 [41, 42]. We constructed two alignments of our 24 newly-generated sequences with these reference sequences using the MAFFT v7.490 plugin in Geneious Prime. The first was an alignment of the complete coding region (CDS), starting with 996 available reference sequences. The second was an alignment of the envelope gene, starting with 1,406 available reference sequences. We then used a subsampling scheme to create smaller datasets amenable to complex inference models [43]. First, we used the jukes-cantor model to create a pairwise genetic distance matrix, and then we selected 30 reference sequences for each of our newly-generated sequences based on closest genetic proximity. We also retained all sequences from Colombia, including those generated during this study as well as publicly available reference sequences. The final DENV-1V datasets contained 209 CDS sequences (83 were from Colombia) and 300 envelope sequences (163 were from Colombia). Finally, to place our DENV1 genotype V sequences in context with other DENV-1 genotypes, we aligned the 209 CDS sequences with 99 representative sequences from DENV-1 genotypes I-V.

### Phylogenetic analysis

We evaluated potential recombination using Phitest and the genetic algorithm for recombination detection (GARD) [44, 45], and did not find any evidence for recombination in either the CDS or envelope dataset.

Maximum likelihood (ML) trees were constructed using IQ-TREE v1.6.12 [46] with the following parameters: 1300 ultrafast bootstraps and 1000 replicates for the SH-like approximate likelihood ratio test. The best-fit model for the CDS dataset was GTR+F+G4, as determined by Bayesian Information Criterion using the ModelFinder software, and the gamma shape alpha was 0.263 [47]. The best fit model for the envelope dataset was TN+F+I+G4, and the gamma shape alpha for the envelope ML tree is 1.292. The ML trees were visualized in iTOL [48] and rooted to the midpoint.

### Molecular clock and phylogeographic inference

To investigate the spatiotemporal dynamics of DENV-1V, we first measured temporal signal using TempEST [49], which evaluates the correlation between sampling dates and genetic distance through root-to-tip regression. Both datasets demonstrated a strong correlation coefficient (>0.92), suggesting sufficient temporal signal for time-scaled phylogenetic analysis using BEAST v2.6.7 [50].

To identify the optimal molecular clock and population growth model, we used a nested sampling (NS) scheme to compare the Strict and Relaxed Log Normal clocks, as well as the Constant, Exponential, Bayesian Skyline and Extended Bayesian Skyline population models using BEAST v2.6.7, BEAUti v2.6.7, and the NS v1.1.0 package. For the envelope data set, a run of 16 particles with a chain length of 20,000 was sufficient to determine the best-fitting model, while for the CDS data set, a total of 60,000 chains with 70 particles were used. The results of these runs were analyzed using NSLogAnalyser in BEAUti v2.6.7.

Phylogeographic reconstruction was performed using the discrete trait model in the BEAST_CLASSIC v1.5.0 package, with the trait location set to department (for Colombian sequences) or country (for non-Colombian sequences), and the optimal molecular clock and population growth models identified above. For the envelope data set, 300 million chains were run, while for the CDS data set, 500 million chains were run. BEAGLE was used to improve run performance [51]. Tracer v1.7.1 [52] was used to visualize parameter distributions and traces for the posteriors, and to verify that ESS values were higher than 300. For each data set, a maximum clade credibility (MCC) tree was summarized using the program TreeAnnotator v2.6.2 with a burn-in percentage of 10. Trees were initially visualized in iTOL [48].

### Selection analysis

We used HyPhy to evaluate both pervasive and episodic selection [53]. We used pervasive selection models SLAC (Single-Likelihood Ancestor Counting), FEL (Fixed Effects likelihood), and FUBAR (Fast, Unconstrained Bayesian AppRoximation), episodic selection model MEME (Mixed Effects Model of Evolution), and aBSREL (adaptive Branch-Site Random Effects Likelihood) model to identify which branches were under selection.

To characterize amino acid changes within our sequences, we reconstructed their MRCA using TreeTime [54]. We identified non-synonymous substitutions compared to this MRCA using Geneious Prime v3.2.1.

## Results

### Molecular testing demonstrates predominance of DENV1 in western Colombia

Between July 2019 and April 2022, a total of 178 febrile patients were recruited in our study site in Risaralda. Among them, 20 dengue infections were confirmed through detection of the NS1 antigen, IgM envelope-specific antibodies, and DENV RNA by RT-qPCR (**Table S2**). This corresponds to 8.2% of the DENV infections reported by the national surveillance system for Risaralda during the same study period. In addition, between August 2021 and November 2022, 201 samples were collected from patients clinically diagnosed with dengue in Valle del Cauca (**Figure S1**). A total of 96 of these samples were positive by the NS1/IgM rapid test, 79 of which were positive for DENV RNA by RT-qPCR (**Table S2**). No infections or coinfections with other arboviruses were identified in patients at either location based on RT-qPCR assay testing for DENV, chikungunya virus, Zika virus, and Mayaro virus.

To identify circulating serotypes, we performed type-specific RT-qPCR. Among the 20 DENV RNA-positive samples from Risaralda, 8 had sufficient DENV RNA concentration for serotype determination; 7 were DENV-1 and 1 was DENV-2. Among the 79 DENV RNA-positive samples from Valle del Cauca, 66 had sufficient DENV RNA for type determination; 42 were DENV-1, 20 were DENV-2, and 4 were DENV-3. Given that DENV-1 was the most prevalent type, we focused our sequencing and phylogenetic analysis on DENV-1. We obtained full-length DENV-1 sequences for seven samples from Risaralda (from 2019-2022) and 17 samples from Valle del Cauca (from 2021-2022) (**Table S3**) using a combination of metagenomic sequencing and multiplex amplicon sequencing.

### Phylogenetic analysis reveals unique regional clades of DENV-1 Genotype V

We sought to define the genetic diversity of DENV-1 in Colombia using our 24 newly-generated sequences as well as publicly available sequences. All of our DENV-1 sequences were classified as genotype V by the DENV genotyping tool [42], and this was confirmed in our initial phylogenetic analysis of all DENV-1 genotypes (**Figure 1A**). According to the new classification proposed by Hill *et al*. [55], 22 of our sequences were classified as Dengue virus serotype 1, genotype V, lineage D.1 (DENV-1V_D.1) and two sequences were classified as Dengue virus serotype 1, genotype V, lineage D.2 (DENV-1V_D.2).

**Figure 1.**
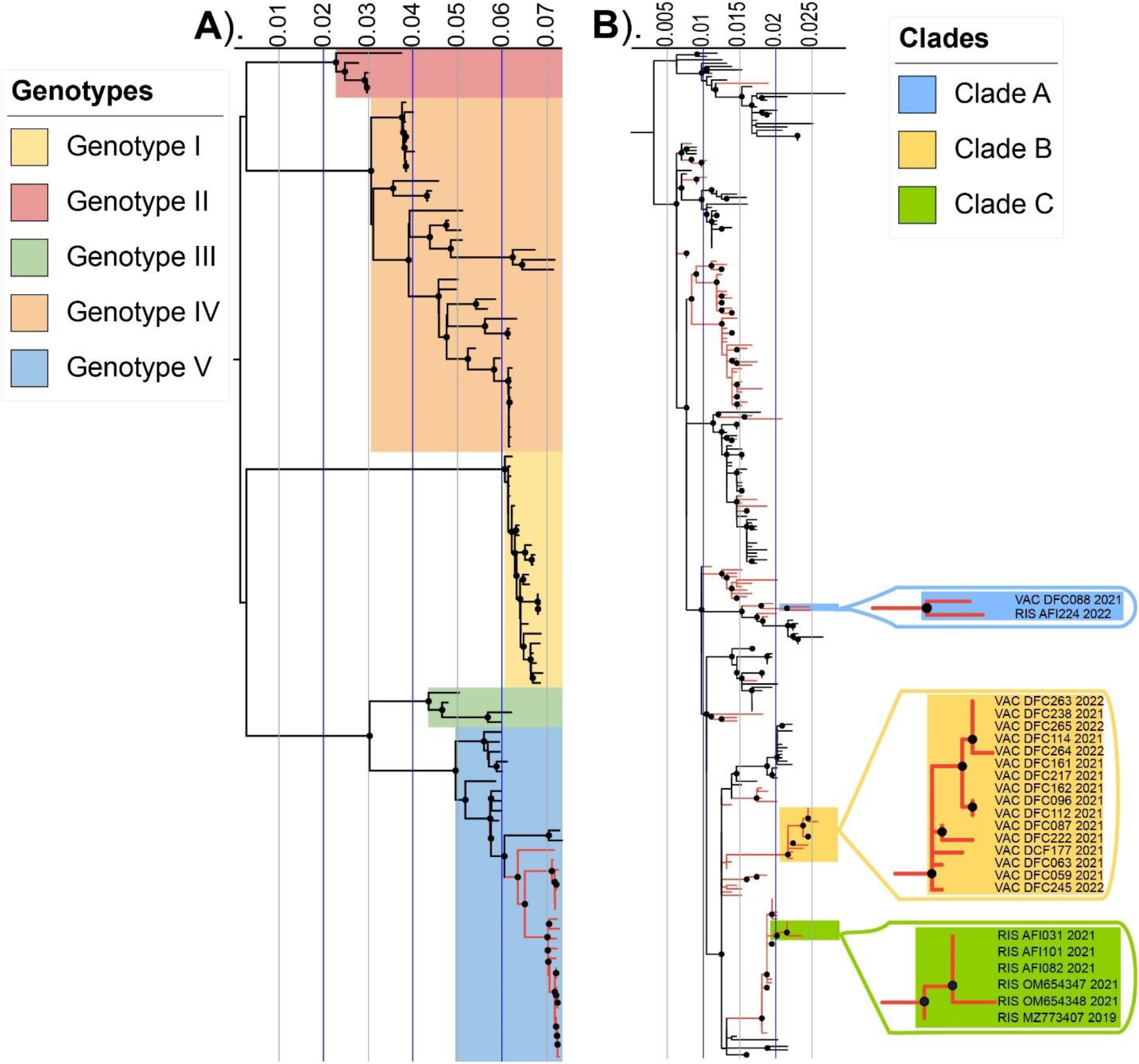
Maximum-likelihood phylogenetic analyses of 24 newly-generated DENV-1 sequences. The black circles on each node represent an ultrafast bootstrap value greater than or equal to 95, and the internal tree scale represents the nucleotide substitutions per site. (A) Full-length representative sequences (n=103) from each DENV-1 genotype, as depicted in the figure legend. The Colombian sequences identified in this study are highlighted in red. (B) Downsampled ML tree of 300 DENV-1 V envelope sequences, including the 24 newly-generated sequences from Colombia. All sequences from Colombia are highlighted in red, and the newly-generated sequences from this study grouped into three distinct clades, as shown in the figure legend.

To further explore their phylogenetic relationships, we constructed an ML tree using all 1,020 available full CDS DENV-1 V reference sequences from NCBI and our 24 new sequences (**Figure S3**) and downsampled this to facilitate further analysis and visualization (**Figure S4**). The resulting phylogenetic trees demonstrated that our sequences clustered into three distinct clades. We repeated this analysis using the *envelope* gene only because there are a larger number of available *envelope* reference sequences (1,406 total, **Figure S5**), downsampled to 300 (**Figure 1B**), capturing greater geographic and temporal diversity than the available full-length sequences.

Analysis of the *envelope* gene confirmed that the sequences generated in this study grouped into three distinct clades, and further demonstrated that each clade was most closely related to other sequences from Colombia (**Figure 1B**). Clade A, within DENV-1V_D.2, contained one sequence from Risaralda in 2022 and one sequence from Valle del Cauca in 2021 (**Figure 1B**, blue highlight), which clustered together with a reference sequence obtained in 2015 from Cundinamarca, a department in central Colombia (**Figure S1**). Clade B contained 16 DENV-1V_D.1 sequences from Valle del Cauca in 2021-2022 (**Figure 1B**, yellow highlight), which clustered with sequences from Santander, a department located in the central northern region of Colombia (**Figure S1**). Clade C contained six DENV-1V_D.1 sequences from Risaralda in 2019-2021 (**Figure 1B**, green highlight), which clustered with sequences obtained in 2016 from Antioquia, an adjacent department located in the northwest of Colombia (**Figure S1**).

Overall, analysis of the *envelope* gene sequences provided more information compared to the full genome sequences, since it allowed us to include closely related viruses for which only the *envelope* gene sequences were available. For example, the relationship between the sequences in Clade B and sequences from Santander would not have been apparent from analysis of only full genome sequences (**Figure S3**). Together, these data demonstrate the circulation of three unique clades of DENV-1V_D circulating at the same time in geographically distinct, but nearby, regions of Colombia.

### Phylodynamic and phylogeographic analyses reveal persistence, evolution, and local transmission of DENV-1 clades in western Colombia

Analyzing the genetic diversity of DENV-1 across temporal and spatial scales can be used to understand how specific lineages are introduced, spread, and maintained in specific regions. To perform phylodynamic analyses, we formally compared molecular clock and population growth models for both the CDS and *envelope* datasets. For the full CDS dataset, the best-fit model used a relaxed lognormal clock with a Bayesian skyline demographic model. For the envelope dataset, a strict molecular clock with a Bayesian skyline demographic model was the best fit, though a relaxed lognormal clock with a Bayesian skyline demographic model had a very similar likelihood (**Table S4**). The most recent common ancestor (MRCA) dates for the two datasets were similar for the nodes of interest, and the confidence intervals were comparable (**Table S5**).

To evaluate the dynamics of DENV-1 V introductions into and across Colombia, we used a discrete phylogeographical model using the *envelope* gene data. Overall, we found that each of Clades A-C was introduced to Risaralda and Valle del Cauca from neighboring departments within Colombia, rather than outside of the country. Our results suggest that these clades and their immediate ancestors have been circulating and evolving within Colombia within the last 15 years. For example, the summarized Maximum Clade Credibility (MCC) tree (**Figure 2A**) estimates the time of the MRCA (tMRCA) of Clade A sequences as January 2018 (HPDI = February 2016 – November 2019, **Figure 2B**). Although the Clade A MRCA was within Colombia, its specific location is uncertain, with a location probability (LP) of 0.37 for Valle del Cauca, 0.29 for Risaralda, and 0.16 for Cundinamarca. The closest relative to Clade A was a sequence obtained from Cundinamarca in 2015. Clade A and this sequence share an MRCA, referred to as A*, that circulated around 2013 (HPDI = February 2012 – February 2015) and whose location was Cundinamarca with a probability of 0.54. Cundinamarca is a department located in the center of Colombia, whose capital is approximately 300 Km from the capital of Risaralda and approximately 450 Km from the capital of Valle del Cauca (**Figure 2C, Figure S1**).

**Figure 2.**
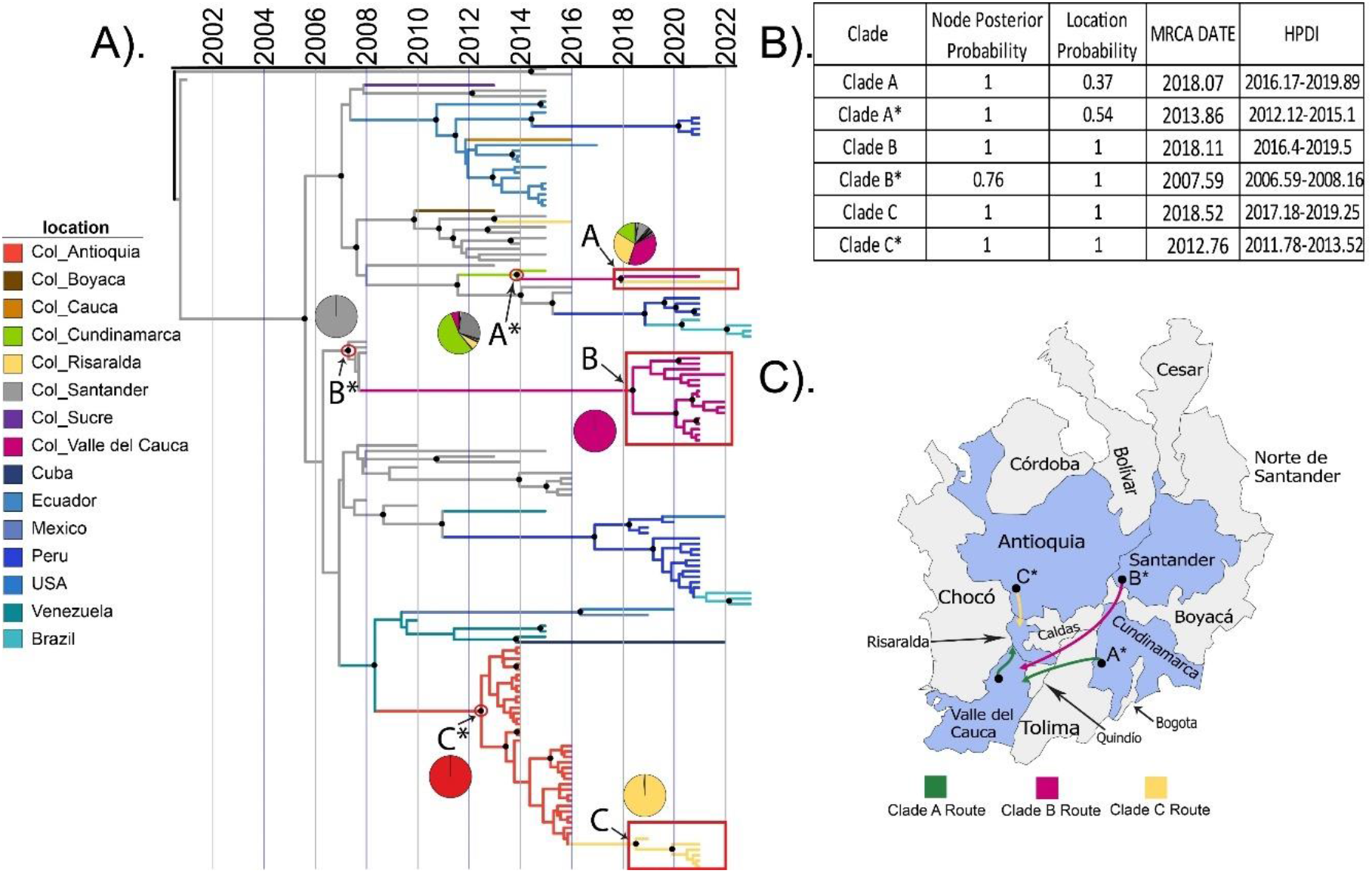
Temporal and geographical analysis of newly-generated DENV sequences. **(A)** Time-scaled Maximum clade credibility (MCC) tree of 147 DENV-1V envelope sequences, including the 24 newly-generated sequences, highlighted with red boxes. Circles indicate nodes with a posterior probability of 90% or greater. Branches are colored by location or inferred location, and pie charts indicate the posterior location probability for each node of interest. Location nomenclature for Colombian sequences includes Col_ and the department name. **(B)** Posterior probability of the node position, location probability and date with 95% highest posterior density (HPD). **(C)** Most probable dispersion routes of clades identified in this study.

The tMRCA of Clade B sequences was February 2018 (HPDI = May 2016 – July 2019). Clade B was most closely related to sequences from Santander, whose MRCA (B*) dates to approximately August 2007 (HPDI = July 2006 - January 2008) with a location in the department of Santander (LP=0.99). The capital of Santander is located 556 Km from the capital of Risaralda and 760 Km from the capital of Valle del Cauca (**Figure S1)**.

The tMRCA of Clade C sequences was July 2018 (HPDI = March 2017 – April 2019). This clade was most closely related to sequences from Antioquia, whose MRCA (C*) dates to October 2012 (HPDI= October 2011 - July 2013) with a location in Antioquia (LP=1.0). The capital of Antioquia is located approximately 250 Km from the capital of Risaralda and approximately 450 Km from the capital of Valle del Cauca (**Figure S1)**.

Comparison of this *envelope* tree and the CDS tree showed similar results and dates for clades A, B, and C (**Figure S6**). Thus, overall, these results indicate that each of the three contemporaneous DENV-1V clades in western Colombia was derived from viruses circulating in nearby regions of Colombia within the last 15 years. Based on this data, the increase in DENV cases observed since 2019 was not due to the introduction of new virus lineages, though the long branches on our phylogenetic tree (especially leading to Clade B) highlight a genomic surveillance gap on DENV circulation and evolution in this region.

Phylogeographic analysis also allowed us to evaluate the dispersion of DENV-1 between Colombia and other countries (**Figure S7**). We observed repeated exchange of DENV-1 between the Colombian departments of Santander and Norte de Santander and the neighboring country of Venezuela until the 2000s (**Figure S6** and **Figure S7**). After the 2000s we observed only two introductions to Colombia, one from Venezuela (LP=0.97) introduced into the department of Antioquia in August 2009 and another from Ecuador (LP= 1), introduced into the department of Cauca in February 2012. On the other hand, two dispersions to other countries with a high probability of ancestral location in Colombia were identified, one to Peru in July 2015 (LP=1) and another to Ecuador in August 2007 (LP=0.99). A third introduction to Venezuela was also identified but in this case with a probability of ancestral location in Colombia of LP=0.46 and a probability of ancestral location in Venezuela of LP=0.34.

### Clade-specific amino acid substitutions occurred across the genome

To evaluate DENV-1 genetic changes that may be associated with recently increased disease burden, we investigated amino acid substitutions and sites under positive selection in contemporary viruses. We identified 25 non-synonymous substitutions in clades A (n=1), B (n=16), and C (n=18) compared to the ancestor of DENV-1V sequences from Colombia (**Table 1**). The highest numbers of substitutions were found in the NS5 (n = 5) and NS1 (n=5) proteins. Clade A had only one clade-specific substitution in the NS3 protein and did not share substitutions with the other clades. Clades B and C shared 12 substitutions with one another. Clade B also had 5 clade-specific substitutions across the prM, envelope, NS2A, NS4B, and NS5 proteins. Clade C had 7 clade-specific substitutions across the prM, envelope, NS1, NS2B, and NS5 protein, though L478I (envelope) and S1405F (NS2B) were only present in sequences from 2021. Thus, DENV-1V viruses circulating contemporaneously in nearby locations were distinct at the amino acid level, including potentially antigenic regions.

**Table 1.**
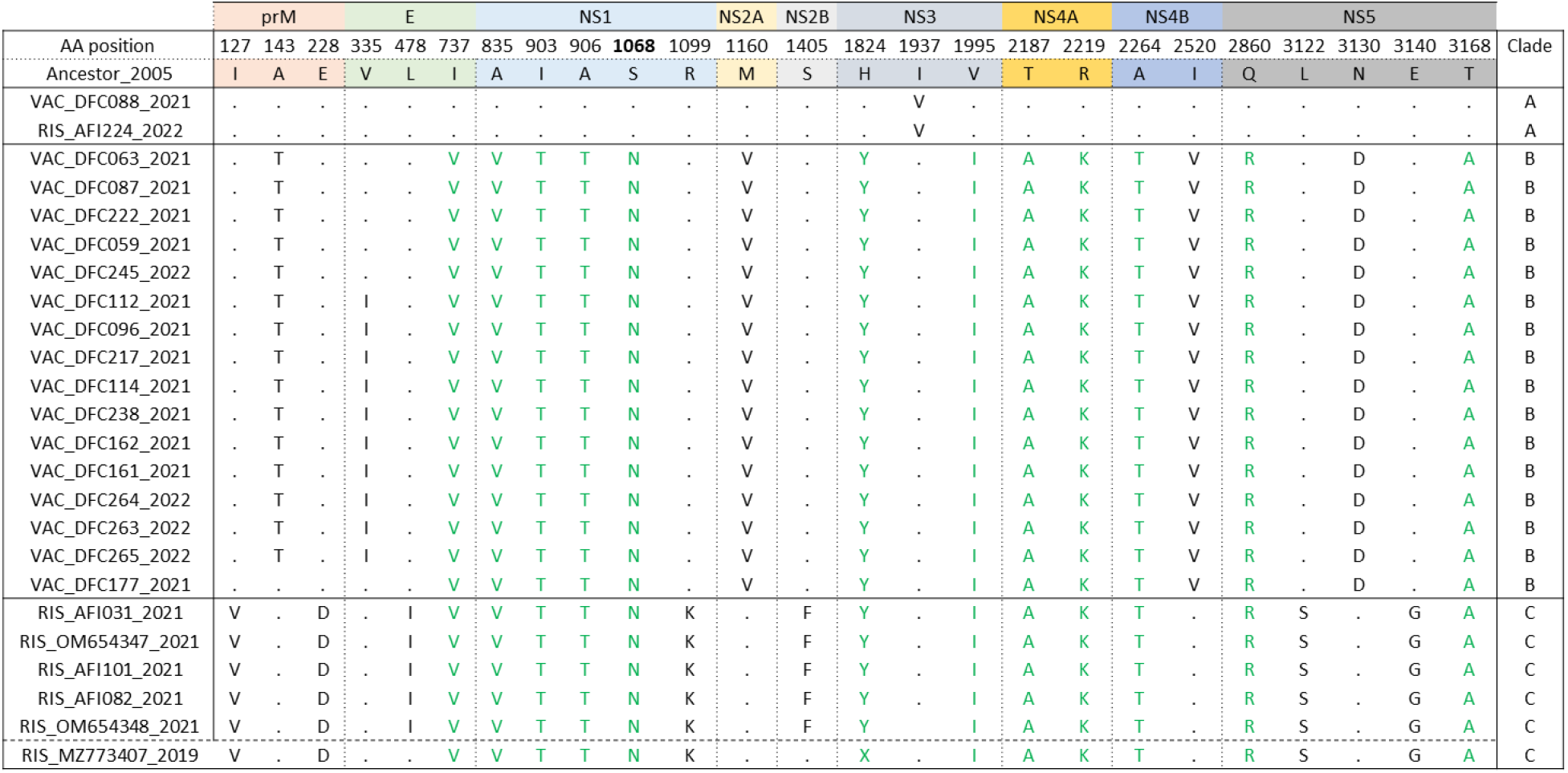
Amino acid changes observed in newly-generated sequences. The substitutions in black are clade-specific, while green represents substitutions found in multiple clades. Each protein region is color-coded, and the dot represents conserved positions. The dashed line separates the Risaralda 2019 sequence from those found in 2021.

We also performed selection analysis using a combination of pervasive and episodic models in HyPhy [53]. From the DENV-1 V sequences represented in our downsampled phylogenetic trees, we identified two codons with a high probability of positive selection (**Table 2**). First, in the CDS dataset, position 293 in the NS1 coding region (corresponding to position 1068 in the full CDS) was identified by MEME, FUBAR, FEL, and SLAC models (p-value <0.05, posterior probability of positive selection/PPPS = 0.99). All branches of Clades B and C were under positive selection at this site, where they shared substitution S293N compared to ancestral sequences. Second, in the envelope dataset, position 248 within the E coding region was identified by FEL, FUBAR, SLAC, and MEME models (p-value <0.07, PPPS = 1). None of the sequences identified in this study had substitutions at this position.

**Table 2.** Positive selection analysis for DENV-1 V sequences from Colombia. Observed amino acid (AA) positions under positive selection in the envelope (n=300) and CDS (n=209) datasets. Pervasive site-based models Fixed Effects Likelihood (FEL), Single-Likelihood Ancestor Counting (SLAC), and Fast, Unconstrained Bayesian AppRoximation (FUBAR) were used to identify positions under persistent selection. The adaptive site-based model, Mixed Effects Model of Evolution (MEME), was used to identify codons under episodic selection. The default settings were used for each model. The position of each codon is relative to the position in each dataset. Selection analysis was implemented using models from HyPhy.

| Dataset | AA<br>Position | Protein | MEME<br>(p-value) | FEL<br>(p-value) | FUBAR<br>(probability) | SLAC<br>(p-value) |
| --- | --- | --- | --- | --- | --- | --- |
| *Envelope* | 248 | E | 0.02 | 0.01 | 1 | 0.07 |
| CDS | 1068 | NS1 | 0.03 | 0.02 | 0.99 | 0.23 |

## Discussion

DENV genomic surveillance is crucial to understanding virus evolution and transmission in endemic locations and during outbreaks. We investigated recent DENV-1 diversity and circulation in Central-West Colombia, a region with very little prior genomic surveillance, during a time of increased disease burden. We detected three distinct clades of DENV-1V: one in Risaralda, one in Valle del Cauca, and one that co-circulated in both departments. We found that each clade had been circulating in nearby departments of Colombia over the preceding 15 years, and thus the increased disease burden observed since 2019 did not reflect introduction of new lineages. Although a number of prior studies have linked DENV outbreaks with clade replacement, there is also increasing evidence that longstanding virus lineages within in a population can contribute to increased disease burden, likely due to shifting environmental factors or demographics [56].

Interestingly, the new 2019-2022 DENV-1 V sequences from Risaralda were not related to the prior DENV-1V sequence reported from that department in 2017, which instead clustered with sequences from Santander. Thus, there appear to have been at least 3 introductions of distinct DENV-1 V clades into Risaralda from different departments to the northeast (Santander, Antioquia, and Valle del Cauca) over the preceding 5 years. We observed similar results for Valle del Cauca, with introductions likely from Santander and Cundinamarca sometime in the last 15 years. Notably, the long branch lengths between our clade MRCAs and their ancestors (especially B and B*) reflect unmeasured circulation and underscore the need for increased genomic surveillance. Because of this, our results must be interpreted with caution, especially since the lack of sequence data from southern and eastern regions of Colombia may mask patterns of transmission and evolution.

On a larger scale, DENV-1V has a wide spatiotemporal dispersion in Colombia and other South American countries and has been the predominant genotype of DENV1 in the Americas since its introduction [25, 57]. DENV1-V was introduced into the Americas from India twice, once in the early 1970s and a second time in the 1990s [24, 58]. The earliest DENV1-V cases were reported in the Caribbean islands and later spread to South America, becoming endemic in many countries [58, 59]. However, in the 1990s, most of the original DENV1-V Caribbean lineages became extinct and were replaced by a lineage that had evolved in Venezuela and another lineage that had evolved in Brazil [25]. Here, we observed repeated exchange of DENV-1V between the Colombian departments of Santander and Norte de Santander and Venezuela, which is consistent with what has been previously reported [20, 57]. We also observed dispersion of DENV-1V from Colombia to Ecuador, consistent with a prior study suggesting that Colombia and Venezuela are the main origins of DENV-1 dissemination to Ecuador [60]. However, this dispersion is not unidirectional since we also identified a strain in the southwestern department of Cauca with common ancestry to clades reported in Ecuador in 2011. Interestingly, sequences recently sampled in Peru (2021) have an MRCA in Colombia with circulation in September 2018, which demonstrates active exchange of DENV between the southwest region of Colombia and neighboring countries. These results highlight Colombia’s role not only as a recipient but also as an exporter of DENV cases.

Future studies and strengthening of genomic surveillance are necessary in these regions to understand how the countries on the southern borders contribute to the exchange of lineages within Colombia.

Similar to our results, prior studies have also demonstrated the importance of local evolution of DENV in Colombia. Laiton-Donato *et al*. reported the divergence of two DENV2 strains with different geographical distributions between 2013-2016 [32]. Salvo *et al*. reported DENV1 clade replacement during an interepidemic season (2014-2016) in a single location (Medellín, Antioquia)[31]. In both examples, virus spread was potentially linked to amino acid substitutions, within the E protein in the first report and NS1 in the second.

Here, we identified clade-specific amino acid mutations in contemporary DENV-1 sequences from Colombia, including 3 mutations in prM. The prM protein forms a heterodimer with the envelope protein, ensuring proper folding and virus particle assembly while preventing premature fusion within intracellular membrane compartments [61]. In a recent study, anti-prM antibodies increased pathogenesis in a mouse model and increased the cell entry of partially mature viruses, which otherwise have reduced entry compared to mature viruses [62, 63]. Thus, substitutions in prM may alter antibody-dependent enhancement. The prM substitution E228D, found in Clade C and its ancestors, is in the highly conserved N-terminus of the a-helical domain. Previous studies have shown that position 228 is involved in prM-E interactions, and we hypothesize that the change from glutamic acid to aspartic acid may alter the binding kinetics for prM and E.

We also identified clade-specific non-synonymous substitutions in the envelope protein. The envelope protein contains three structural domains (DI-DIII) and a transmembrane anchor, which is involved in targeting it to sites of virion assembly [64]. D-I organizes the structure of mature dimers, D-II consists of the dimerization and fusion loop, and D-III consists of the Ig-like receptor-binding region involved in viral entry [65, 66]. In Clade C, we identified a DII substitution at L478I in sequences from 2021 but not 2019. Interestingly, this mutation is predicted to increase hydrophobicity, and previous studies have shown that substitutions that increased hydrophobicity in DII contribute to immune escape [67]. Finally, Clades C and B shared a common substitution (I737V) with sequences from Santander in 2001. This position is in the first α-helix of the transmembrane domain, which is primarily involved in the assembly of virus particles, so substitutions here may affect virion assembly [68]. Further investigation is needed to characterize the potential functional relevance of these substitutions to pathogenesis and immune evasion.

In addition to these mutations in structural proteins, we also identified several non-synonymous mutations in NS1, including one site (S1068N) under positive selection. Interestingly, Salvo *et al*. also reported positive selection in NS1 among DENV-1 sequences from Colombia, though not at this specific position [31]. NS1 is a multi-function protein with two states: a membrane-associated dimer involved in intracellular viral replication, and an oligomeric secreted form. The secreted form is linked to increased viral pathogenesis through the induction of endothelial dysfunction, vascular leak, and pro-inflammatory cytokine release from immune cells [69-73]. The NS1 protein comprises the N-terminal B-roll, Wing (surrounded by connector domains), and β-Ladder domain [74]. Previous studies have shown that antibodies targeting either the wing or β-ladder domains can block NS1 binding to endothelial or immune cells and are protective in lethal challenge models [69, 75-83]. Recently, highly protective NS1 antibodies have been mapped to conserved regions in the NS1 β-ladder domain that are directly adjacent to the S1068N mutation we identified (R1074 and T1076) [80]. Therefore, we hypothesize that this substitution in the β-ladder may be involved in immune escape through glycosylation alterations or steric hindrance. We also identified three shared substitutions between Clades B and C in the Wing domain, including threonine, serine, or asparagine substitutions that could support glycosylation or polarity changes that alter immune recognition [84]. Further work is needed to assess the role of these mutations in pathogenesis and immune evasion.

Overall, analysis of the sequences generated in this study revealed that contemporary DENV-1V viruses descended from viruses that have been circulating in Colombia over the last 15 years. Despite recent increases in disease burden, we did not find evidence for the introduction of new lineages or sub-lineages. We also observed co-circulation of multiple DENV-1V clades in nearby geographic regions, as well as clade-specific mutations, some of which may have a role in immune evasion or pathogenesis. Further work is needed to understand DENV evolution and dispersion in the context of the local immune landscape.

## Author Contributions

DRG, MHC, JACO, and AP conceived of and designed the study; DRG, BG, DL, PE, EJP, JRC, MMG, JRS, JO, and JJW generated and analyzed epidemiological and diagnostic data; TF, AK, DK, NS, MAM and AB generated and analyzed sequence data; DRG, TF, AB, and AP performed phylogenetic and phylogeographic analysis; DRG, TF, CJN and AP drafted the manuscript; all authors reviewed and edited the manuscript.

## Acknowledgements

We thank Dr. Daniel Espinoza for logistical support.

## Data availability

The sequences generated in this study have been deposited in the NCBI (National Center for Biotechnology Information) GenBank database. These sequences are publicly available to the scientific community under the accessions reported in supplementary material table S3. In addition, the corresponding Sequence Read Archive (SRA) has been uploaded and is accessible in the NCBI SRA database. These resources allow the reproduction of the analyses carried out and facilitate future studies.

## References

1. Guzmán, M.G. and G. Kouri, Dengue: an update. The Lancet infectious diseases, 2002. 2(1): p. 33–42.

2. Bhatt, S., et al., The global distribution and burden of dengue. Nature, 2013. 496(7446): p. 504–507.

3. Urdaneta-Marquez, L. and A.-B. Failloux, Population genetic structure of Aedes aegypti, the principal vector of dengue viruses. Infection, Genetics and Evolution, 2011. 11(2): p. 253–261.

4. Rezza, G., Aedes albopictus and the reemergence of Dengue. BMC public health, 2012. 12: p. 1–3.

5. Muñoz, E., et al., Spatiotemporal dynamics of dengue in Colombia in relation to the combined effects of local climate and ENSO. Acta Tropica, 2021. 224: p. 106136.

6. Ordonez-Sierra, G., et al., Multilevel analysis of social, climatic and entomological factors that influenced dengue occurrence in three municipalities in Colombia. One Health, 2021. 12: p. 100234.

7. Alied, M., et al., Latin America in the clutches of an old foe: Dengue. Brazilian Journal of Infectious Diseases, 2023. 27: p. 102788.

8. Gutierrez, L. PAHO/WHO data - incidence: PAHO/WHO. Pan American Health Organization 2015 [cited 2024 03/07/2024]; Available from: https://www3.paho.org/data/index.php/en/mnu-topics/indicadores-dengue-en/dengue-regional-en/315-reg-dengue-incidence-en.html.

9. PAHO/WHO Situation report no 9 - dengue epidemiological situation in the region of the Americas - Epidemiological week 08, 2024. Pan American Health Organization 2024 [cited 2024 03/20/2024]; Available from: https://www.paho.org/en/documents/situation-report-no-9-dengue-epidemiological-situation-region-americas-epidemiological.

10. Jing, Q. and M. Wang, Dengue epidemiology. Global Health Journal, 2019. 3(2): p. 37–45.

11. Guzman, M.G., et al., Dengue: a continuing global threat. Nature reviews microbiology, 2010. 8(Suppl 12): p. S7–S16.

12. Thongsripong, P., et al., Phylodynamics of dengue virus 2 in Nicaragua leading up to the 2019 epidemic reveals a role for lineage turnover. BMC Ecology and Evolution, 2023. 23(1): p. 58.

13. Kuno, G., et al., Phylogeny of the genus Flavivirus. Journal of virology, 1998. 72(1): p. 73–83.

14. Poltep, K., et al., Genetic diversity of dengue virus in clinical specimens from Bangkok, Thailand, during 2018–2020: Co-circulation of all four serotypes with multiple genotypes and/or clades. Tropical medicine and infectious disease, 2021. 6(3): p. 162.

15. Gallichotte, E.N., et al., Genetic variation between dengue virus type 4 strains impacts human antibody binding and neutralization. Cell reports, 2018. 25(5): p. 1214–1224.

16. Weaver, S.C. and N. Vasilakis, Molecular evolution of dengue viruses: contributions of phylogenetics to understanding the history and epidemiology of the preeminent arboviral disease. Infection, genetics and evolution, 2009. 9(4): p. 523–540.

17. Yenamandra, S.P., et al., Evolution, heterogeneity and global dispersal of cosmopolitan genotype of Dengue virus type 2. Scientific Reports, 2021. 11(1): p. 13496.

18. Rico-Hesse, R., Molecular evolution and distribution of dengue viruses type 1 and 2 in nature. Virology, 1990. 174(2): p. 479–493.

19. Boshell, J., et al., Dengue en Colombia. Biomédica, 1986. 6(3-4): p. 101–106.

20. Carrillo-Hernandez, M.Y., et al., Phylogenetic and evolutionary analysis of dengue virus serotypes circulating at the Colombian–Venezuelan border during 2015–2016 and 2018–2019. PLoS One, 2021. 16(5): p. e0252379.

21. Padilla, J., D. Rojas, and R. Sáenz-Gómez, Dengue in Colombia: Epidemiology of Hyperendemic Reemergence. Guías de impresión (Bogotá, Colombia, 2012.

22. Gutierrez-Barbosa, H., et al., Dengue infections in Colombia: epidemiological trends of a hyperendemic country. Tropical medicine and infectious disease, 2020. 5(4): p. 156.

23. Cáceres, C., et al., Complete nucleotide sequence analysis of a Dengue-1 virus isolated on Easter Island, Chile. Archives of virology, 2008. 153: p. 1967–1970.

24. Villabona-Arenas, C.J. and P.M.d.A. Zanotto, Worldwide spread of dengue virus type 1. PloS one, 2013. 8(5): p. e62649.

25. de Bruycker-Nogueira, F., et al., Evolutionary history and spatiotemporal dynamics of DENV-1 genotype V in the Americas. Infection, Genetics and Evolution, 2016. 45: p. 454–460.

26. Ramos-Castañeda, J., et al., Dengue in Latin America: systematic review of molecular epidemiological trends. PLoS neglected tropical diseases, 2017. 11(1): p. e0005224.

27. Reyes, A., D. Ruge, and L. Herrera, Informe de evento Dengue, Colombia, 2019. Instituto Nacional de Salud, Colombia [Internet], 2019.

28. O’Connor, O., et al., Potential role of vector-mediated natural selection in dengue virus genotype/lineage replacements in two epidemiologically contrasted settings. Emerging microbes & infections, 2021. 10(1): p. 1346–1357.

29. Lambrechts, L., et al., Dengue-1 virus clade replacement in Thailand associated with enhanced mosquito transmission. Journal of virology, 2012. 86(3): p. 1853–1861.

30. OhAinle, M., et al., Dynamics of dengue disease severity determined by the interplay between viral genetics and serotype-specific immunity. Science translational medicine, 2011. 3(114): p. 114ra128–114ra128.

31. Salvo, M.A., et al., Tracking dengue virus type 1 genetic diversity during lineage replacement in an hyperendemic area in Colombia. Plos one, 2019. 14(3): p. e0212947.

32. Laiton-Donato, K., et al., Molecular characterization of dengue virus reveals regional diversification of serotype 2 in Colombia. Virology journal, 2019. 16: p. 1–7.

33. Waggoner, J.J., et al., Single-reaction multiplex reverse transcription PCR for detection of Zika, chikungunya, and dengue viruses. Emerging infectious diseases, 2016. 22(7): p. 1295.

34. Waggoner, J.J., et al., Single-reaction, multiplex, real-time rt-PCR for the detection, quantitation, and serotyping of dengue viruses. PLoS neglected tropical diseases, 2013. 7(4): p. e2116.

35. Langsjoen, R.M., et al., Eastern Equine Encephalitis Virus Diversity in Massachusetts Patients, 1938–2020. The American Journal of Tropical Medicine and Hygiene, 2023. 109(2): p. 387.

36. Park, D., broadinstitute/viral-ngs: v1. 25.0.(2019). URL 10.5281/zenodo.3509008.

37. Su, W., et al., A Serotype-Specific and Multiplex PCR Method for Whole-Genome Sequencing of Dengue Virus Directly from Clinical Samples. Microbiology Spectrum, 2022. 10(5): p. e01210–22.

38. Quick, J., et al., Multiplex PCR method for MinION and Illumina sequencing of Zika and other virus genomes directly from clinical samples. Nature protocols, 2017. 12(6): p. 1261–1276.

39. Glenn, T.C., et al., Adapterama II: universal amplicon sequencing on Illumina platforms (TaggiMatrix). PeerJ, 2019. 7: p. e7786.

40. Patel, H., nf-core/viralrecon: nf-core/viralrecon v2. 6.0-Rhodium Raccoon (2023).

41. Fonseca, V., et al., A computational method for the identification of Dengue, Zika and Chikungunya virus species and genotypes. PLoS neglected tropical diseases, 2019. 13(5): p. e0007231.

42. Vilsker, M., et al., Genome Detective: an automated system for virus identification from high-throughput sequencing data. Bioinformatics, 2019. 35(5): p. 871–873.

43. Mongiardino Koch, N., Phylogenomic subsampling and the search for phylogenetically reliable loci. Molecular Biology and Evolution, 2021. 38(9): p. 4025–4038.

44. Huson, D.H., T. Kloepper, and D. Bryant, SplitsTree 4.0-Computation of phylogenetic trees and networks. Bioinformatics, 2008. 14: p. 68–73.

45. Kosakovsky Pond, S.L., et al., GARD: a genetic algorithm for recombination detection. Bioinformatics, 2006. 22(24): p. 3096–3098.

46. Nguyen, L.-T., et al., IQ-TREE: a fast and effective stochastic algorithm for estimating maximum-likelihood phylogenies. Molecular biology and evolution, 2015. 32(1): p. 268–274.

47. Kalyaanamoorthy, S., et al., ModelFinder: fast model selection for accurate phylogenetic estimates. Nature methods, 2017. 14(6): p. 587–589.

48. Letunic, I. and P. Bork, Interactive Tree Of Life (iTOL) v5: an online tool for phylogenetic tree display and annotation. Nucleic acids research, 2021. 49(W1): p. W293–W296.

49. Rambaut, A., et al., Exploring the temporal structure of heterochronous sequences using TempEst (formerly Path-O-Gen). Virus evolution, 2016. 2(1): p. vew007.

50. Bouckaert, R., et al., BEAST 2.5: An advanced software platform for Bayesian evolutionary analysis. PLoS computational biology, 2019. 15(4): p. e1006650.

51. Ayres, D.L., et al., BEAGLE: an application programming interface and high-performance computing library for statistical phylogenetics. Systematic biology, 2012. 61(1): p. 170–173.

52. Rambaut, A., et al., Posterior summarization in Bayesian phylogenetics using Tracer 1.7. Systematic biology, 2018. 67(5): p. 901–904.

53. Kosakovsky Pond, S.L., et al., HyPhy 2.5—a customizable platform for evolutionary hypothesis testing using phylogenies. Molecular biology and evolution, 2020. 37(1): p. 295–299.

54. Sagulenko, P., V. Puller, and R.A. Neher, TreeTime: Maximum-likelihood phylodynamic analysis. Virus evolution, 2018. 4(1): p. vex042.

55. Hill, V., et al., A new lineage nomenclature to aid genomic surveillance of dengue virus. medRxiv, 2024.

56. Ashall, J., et al., A phylogenetic study of dengue virus in urban Vietnam shows long-term persistence of endemic strains. Virus evolution, 2023. 9(1): p. vead012.

57. Jiménez-Silva, C.L., et al., Evolutionary history and spatio-temporal dynamics of dengue virus serotypes in an endemic region of Colombia. PloS one, 2018. 13(8): p. e0203090.

58. Walimbe, A.M., et al., Global phylogeography of Dengue type 1 and 2 viruses reveals the role of India. Infection, genetics and evolution, 2014. 22: p. 30–39.

59. Ribeiro, G.d.O., et al., Adaptive evolution of new variants of dengue virus serotype 1 genotype V circulating in the Brazilian Amazon. Viruses, 2021. 13(4): p. 689.

60. Maljkovic Berry, I., et al., The origins of dengue and chikungunya viruses in Ecuador following increased migration from Venezuela and Colombia. BMC Evolutionary Biology, 2020. 20: p. 1–12.

61. Wang, P.-G., et al., Efficient assembly and secretion of recombinant subviral particles of the four dengue serotypes using native prM and E proteins. PLoS One, 2009. 4(12): p. e8325.

62. Dejnirattisai, W., et al., Cross-reacting antibodies enhance dengue virus infection in humans. Science, 2010. 328(5979): p. 745–748.

63. Dowd, K.A., et al., prM-reactive antibodies reveal a role for partially mature virions in dengue virus pathogenesis. Proceedings of the National Academy of Sciences, 2023. 120(3): p. e2218899120.

64. Klein, D.E., J.L. Choi, and S.C. Harrison, Structure of a dengue virus envelope protein late-stage fusion intermediate. Journal of virology, 2013. 87(4): p. 2287–2293.

65. Modis, Y., et al., Structure of the dengue virus envelope protein after membrane fusion. Nature, 2004. 427(6972): p. 313–319.

66. Kuhn, R.J., et al., Structure of dengue virus: implications for flavivirus organization, maturation, and fusion. Cell, 2002. 108(5): p. 717–725.

67. Messer, W.B., et al., Dengue virus envelope protein domain I/II hinge determines long-lived serotype-specific dengue immunity. Proceedings of the National Academy of Sciences, 2014. 111(5): p. 1939–1944.

68. Hsieh, S.-C., W.-Y. Tsai, and W.-K. Wang, The length of and nonhydrophobic residues in the transmembrane domain of dengue virus envelope protein are critical for its retention and assembly in the endoplasmic reticulum. Journal of virology, 2010. 84(9): p. 4782–4797.

69. Beatty, P.R., et al., Dengue virus NS1 triggers endothelial permeability and vascular leak that is prevented by NS1 vaccination. Sci Transl Med, 2015. 7(304): p. 304ra141.

70. Modhiran, N., et al., Dengue virus NS1 protein activates cells via Toll-like receptor 4 and disrupts endothelial cell monolayer integrity. Sci Transl Med, 2015. 7(304): p. 304ra142.

71. Glasner, D.R., et al., Dengue virus NS1 cytokine-independent vascular leak is dependent on endothelial glycocalyx components. PLoS Pathog, 2017. 13(11): p. e1006673.

72. Puerta-Guardo, H., et al., Flavivirus NS1 triggers tissue-specific vascular endothelial dysfunction reflecting disease tropism. Cell Rep, 2019. 26(6): p. 1598-1613.e8.

73. Valero, N., et al., Differential induction of cytokines by human neonatal, adult, and elderly monocyte/macrophages infected with dengue virus. Viral immunology, 2014. 27(4): p. 151–159.

74. Akey, D.L., et al., Flavivirus NS1 structures reveal surfaces for associations with membranes and the immune system. Science, 2014. 343(6173): p. 881–885.

75. Brault, A.C., et al., A Zika Vaccine Targeting NS1 Protein Protects Immunocompetent Adult Mice in a Lethal Challenge Model. Sci Rep, 2017. 7(1): p. 14769.

76. Bailey, M.J., et al., Antibodies elicited by an NS1-based vaccine protect mice against Zika virus. mBio, 2019. 10(2): p. 02861–18.

77. Espinosa, D.A., et al., Cyclic dinucleotide-adjuvanted dengue virus nonstructural protein 1 induces protective antibody and T cell responses. J Immunol, 2019. 202(4): p. 1153–1162.

78. Lai, Y.C., et al., Antibodies against modified NS1 wing domain peptide protect against dengue virus infection. Sci Rep, 2017. 7(1): p. 6975.

79. Bailey, M.J., et al., Human antibodies targeting Zika virus NS1 provide protection against disease in a mouse model. Nat Commun, 2018. 9(1): p. 4560.

80. Biering, S.B., et al., Structural basis for antibody inhibition of flavivirus NS1-triggered endothelial dysfunction. Science, 2021. 371(6525): p. 194–200.

81. Wessel, A.W., et al., Human monoclonal antibodies against NS1 protein protect against lethal West Nile virus infection. mBio, 2021. 12(5): p. e0244021.

82. Modhiran, N., et al., A broadly protective antibody that targets the flavivirus NS1 protein. Science, 2021. 371(6525): p. 190–194.

83. Lai, Y.-C., et al., Antibodies against modified NS1 wing domain peptide protect against dengue virus infection. Scientific reports, 2017. 7(1): p. 6975.

84. Tan, B.E., M.R. Beard, and N.S. Eyre, Identification of Key Residues in Dengue Virus NS1 Protein That Are Essential for Its Secretion. Viruses, 2023. 15(5): p. 1102.

