## Supplemental Material for "Spatiotemporal dispersion of DENV1 genotype V in western Colombia"

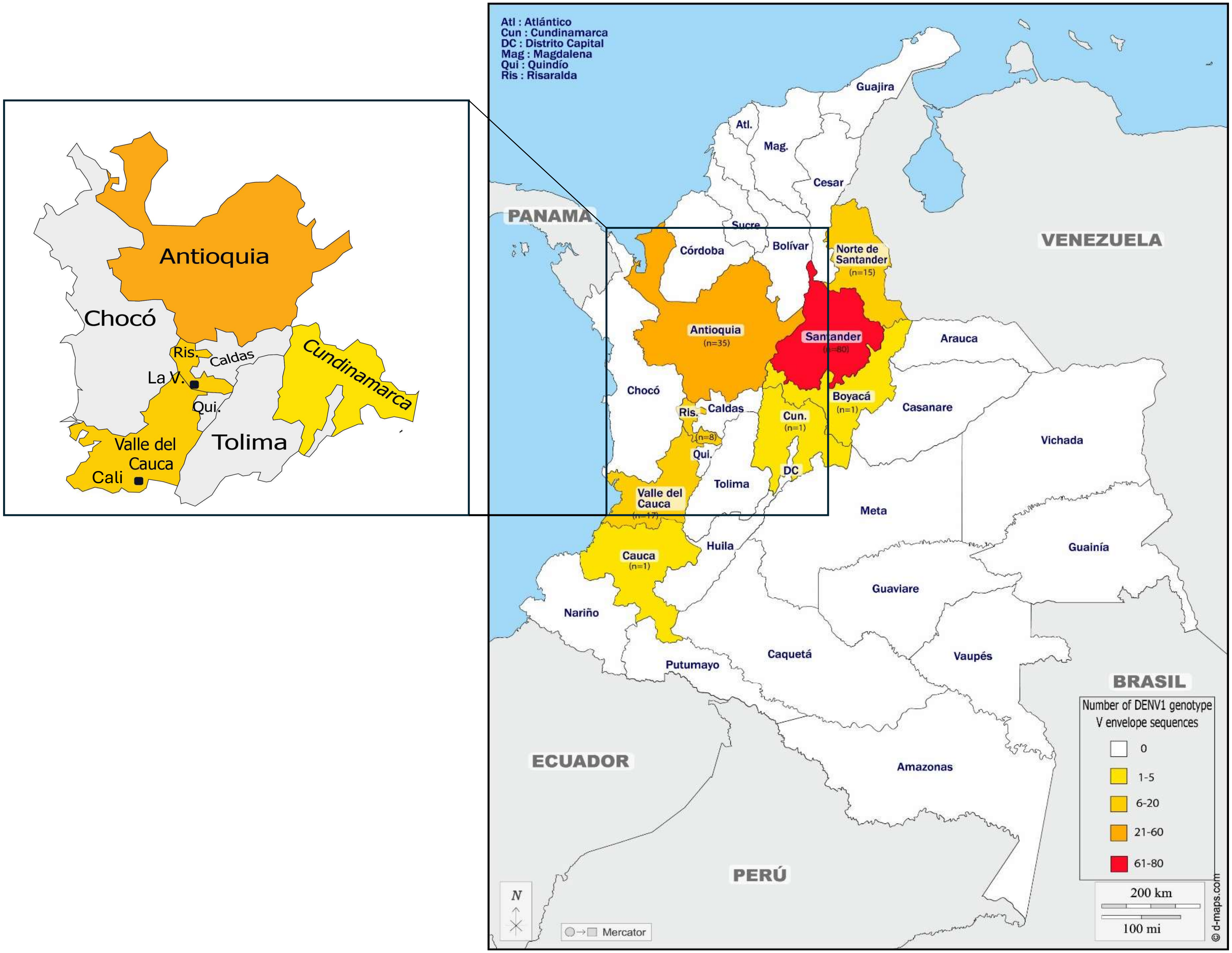


**Figure S1. Departmental map of Colombia.** Departments are colored by the number of DENV1 genotype V envelope genome sequences available for analysis, including those generated in this study. The departments of Antioquia, Cundinamarca, Norte de Santander, Risaralda, and Santander belong to the Andean region, while the departments of Cauca and Valle del Cauca belong to the Pacific region, and the department of Sucre belongs to the Caribbean region. The inset shows the two cities included in this study: La Virginia (La V.) and Cali.

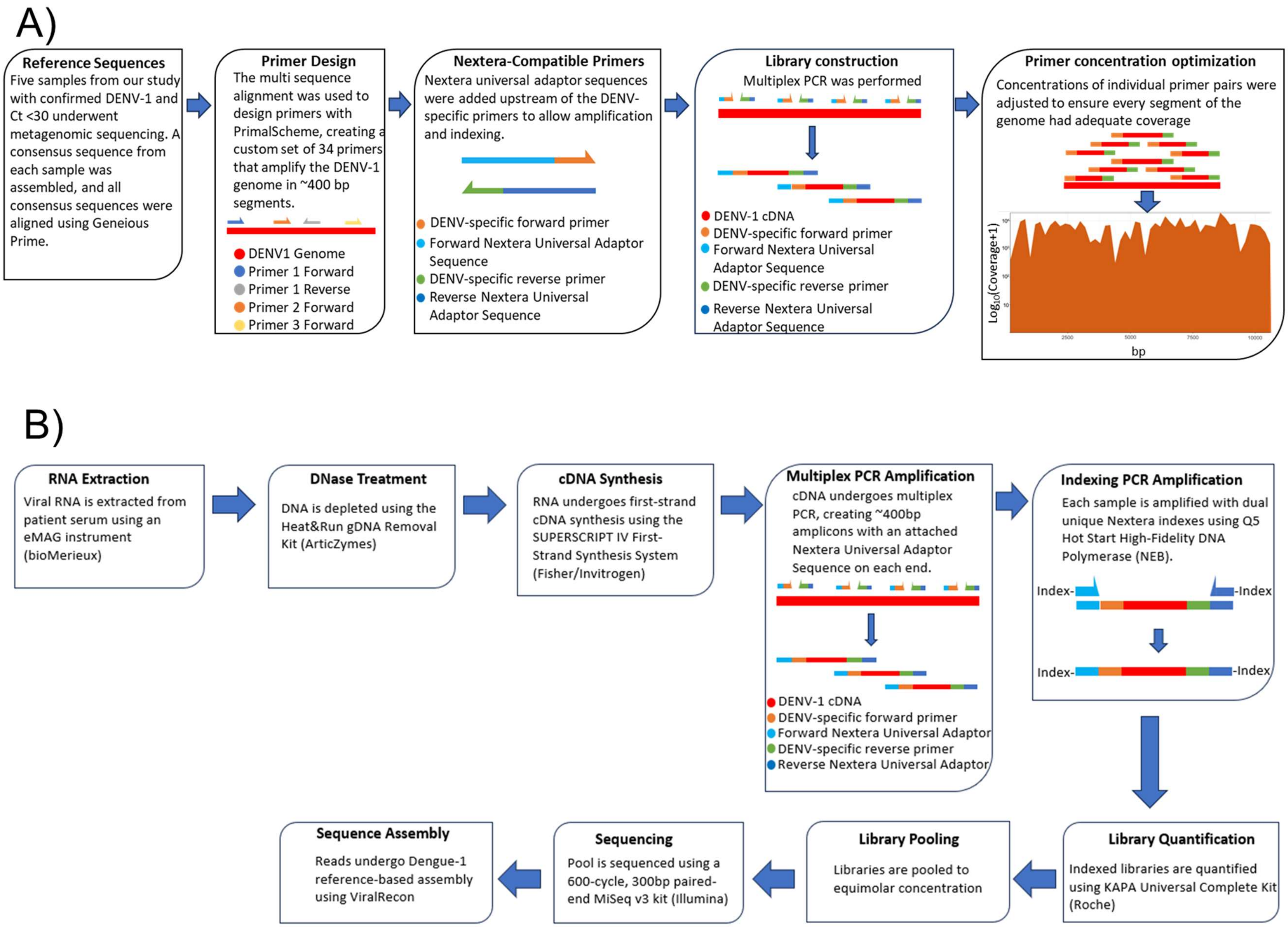


**Figure S2. Amplicon-based sequencing approach.** (A) Steps used to create custom primers for multiplex amplicon sequencing of DENV1 genotype V in this study. (B) Laboratory and analysis workflow.

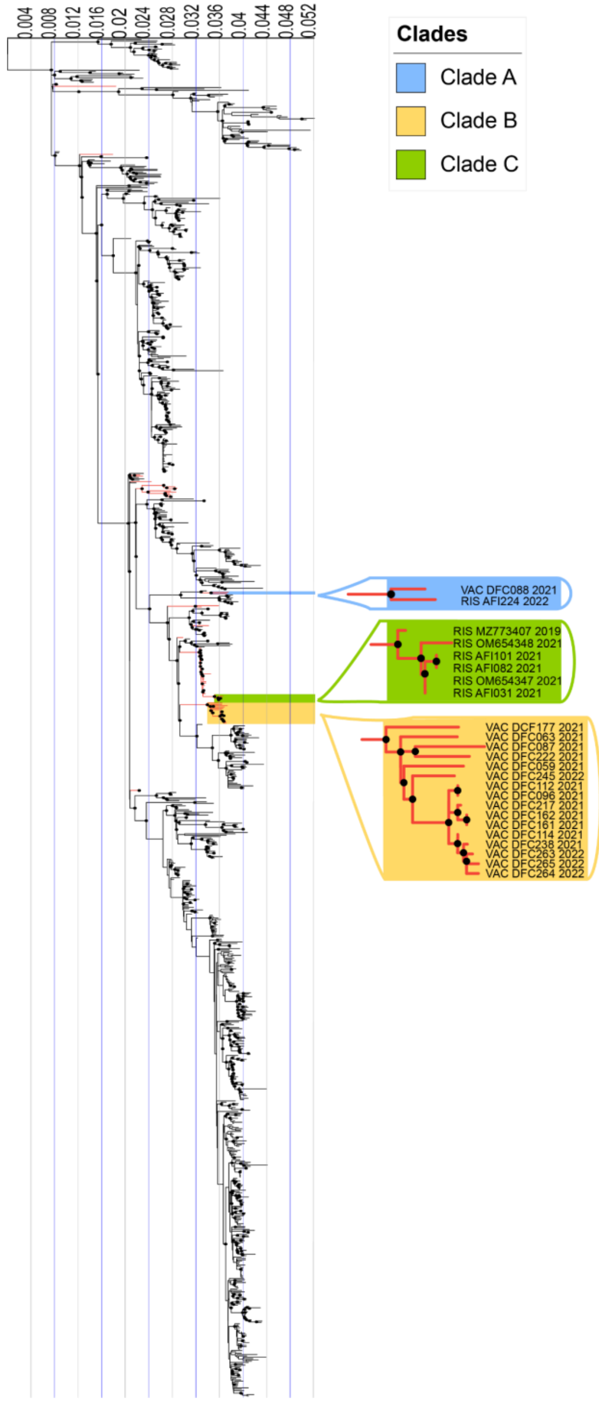


**Figure S3**. **Maximum-likelihood tree of 1,020 full-length DENV1 genotype V sequences**. Twenty-four newly-generated sequences from Colombia are highlighted with red branches and fall into three distinct clades, as shown in the legend. The black circles on each node represent an ultrafast bootstrap value greater than or equal to 95, and the internal tree scale represents the nucleotide substitution per site.

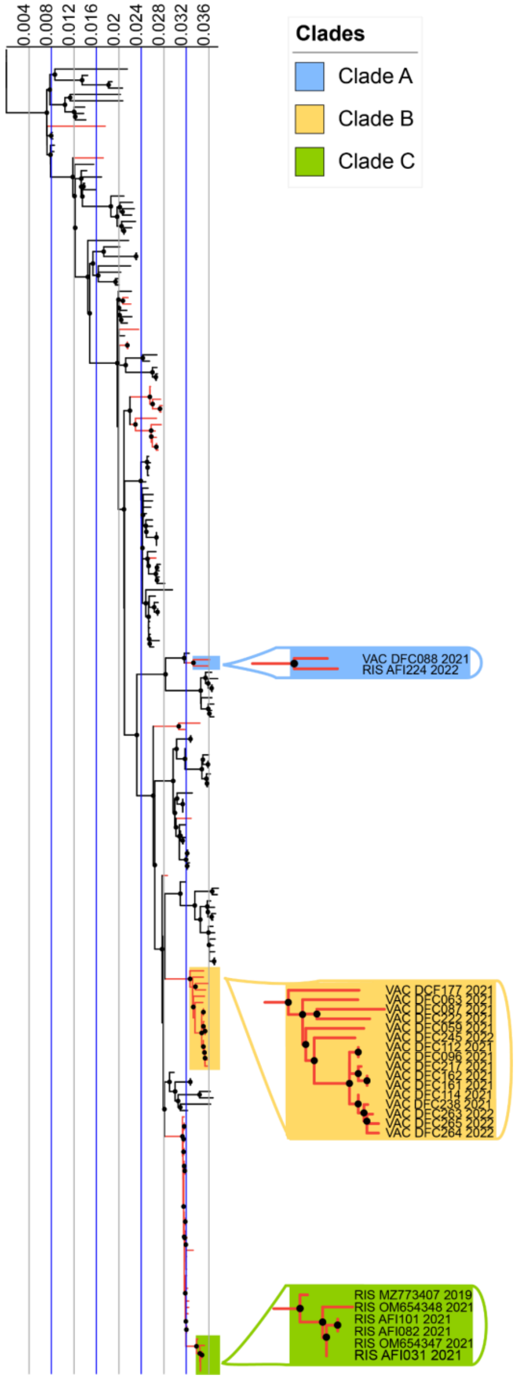


**Figure S4. Downsampled maximum-likelihood tree of 209 full-length DENV1 genotype V sequences.** Twenty-four newly-generated sequences from Colombia are highlighted with red branches and fall into three distinct clades, as shown in the legend. The black circles on each node represent an ultrafast bootstrap value greater than or equal to 95, and the internal tree scale represents the nucleotide substitution per site.

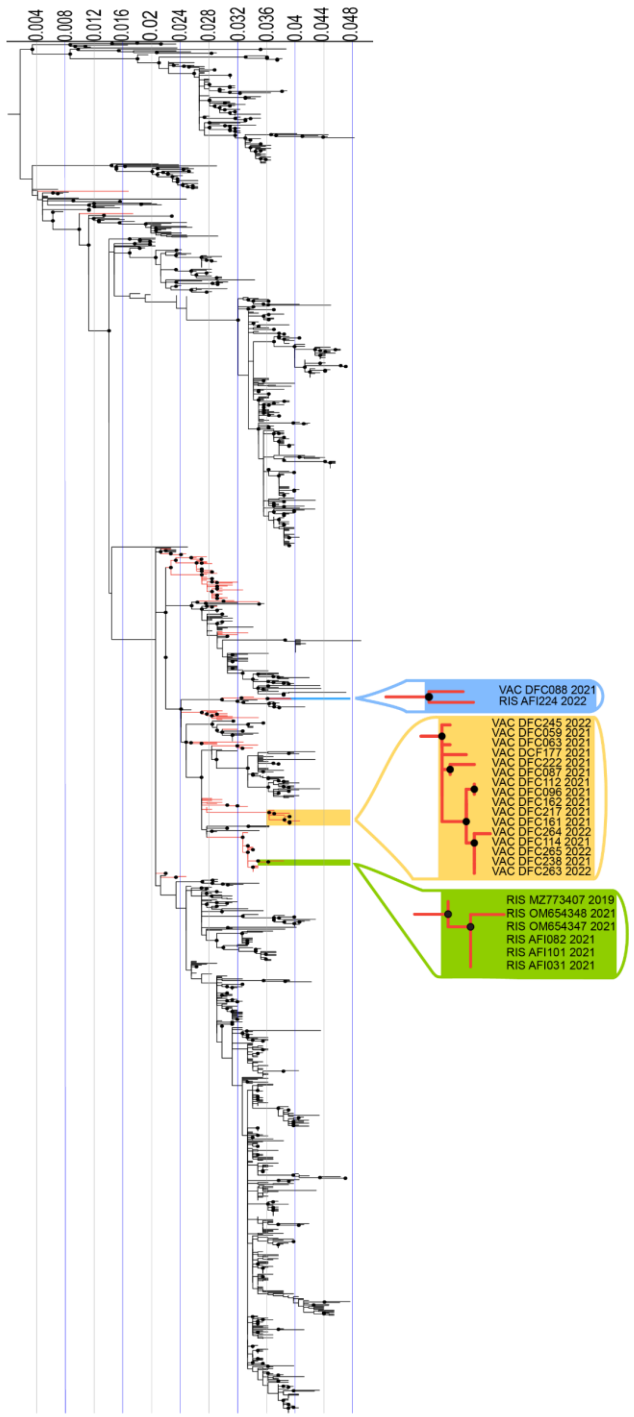


**Figure S5**. **Maximum-likelihood tree of 1,406 envelope DENV1 genotype V sequences**. Twenty-four newly-generated sequences from Colombia are highlighted with red branches and fall into three distinct clades, as shown in the legend. The black circles on each node represent an ultrafast bootstrap value greater than or equal to 95, and the internal tree scale represents the nucleotide substitution per site.

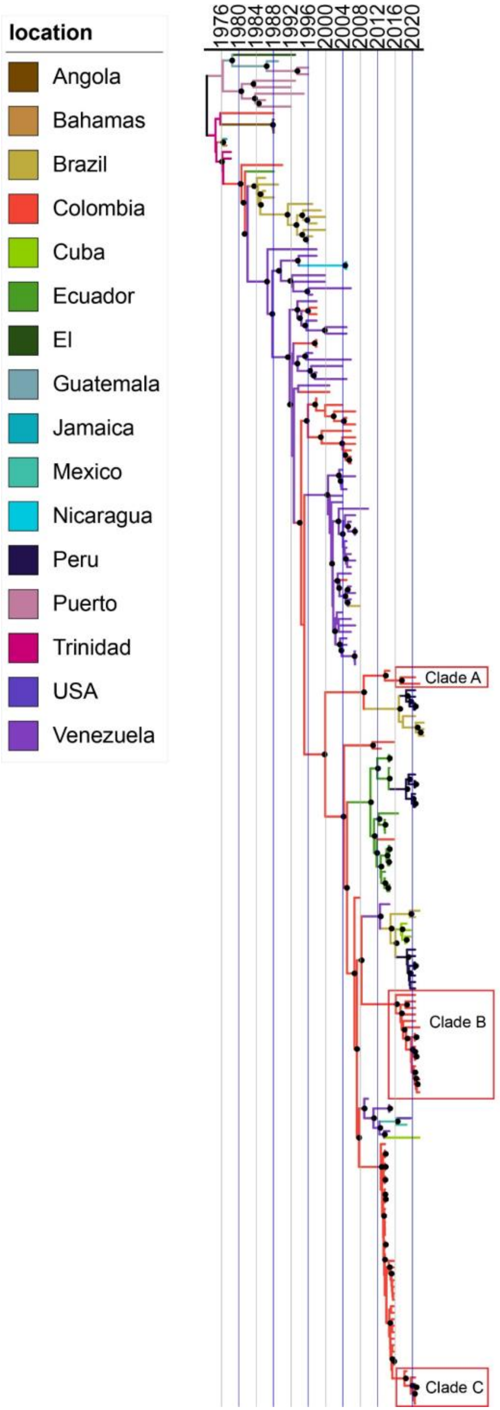


| Clade | Node posterior Probability | Location probability | TMRCA | 95% HPD |
|---|---|---|---|---|
| A | 1 | 1 | 2017.82 | 2015.73-2019.84 |
| B | 1 | 1 | 2016.69 | 2015.01-2018.38 |
| C | 1 | 1 | 2018.69 | 2017.95-2019.21 |

**Figure S6**. Time-scaled Maximum clade credibility (MCC) tree of 209 full-length DENV1 genotype V CDS sequences, including 24 newly-generated sequences. Sequences highlighted in red were identified in this study, while other sequences were obtained from GenBank. Branch colors are based on the location of the isolated virus samples and inferred probable posterior location of its ancestors.

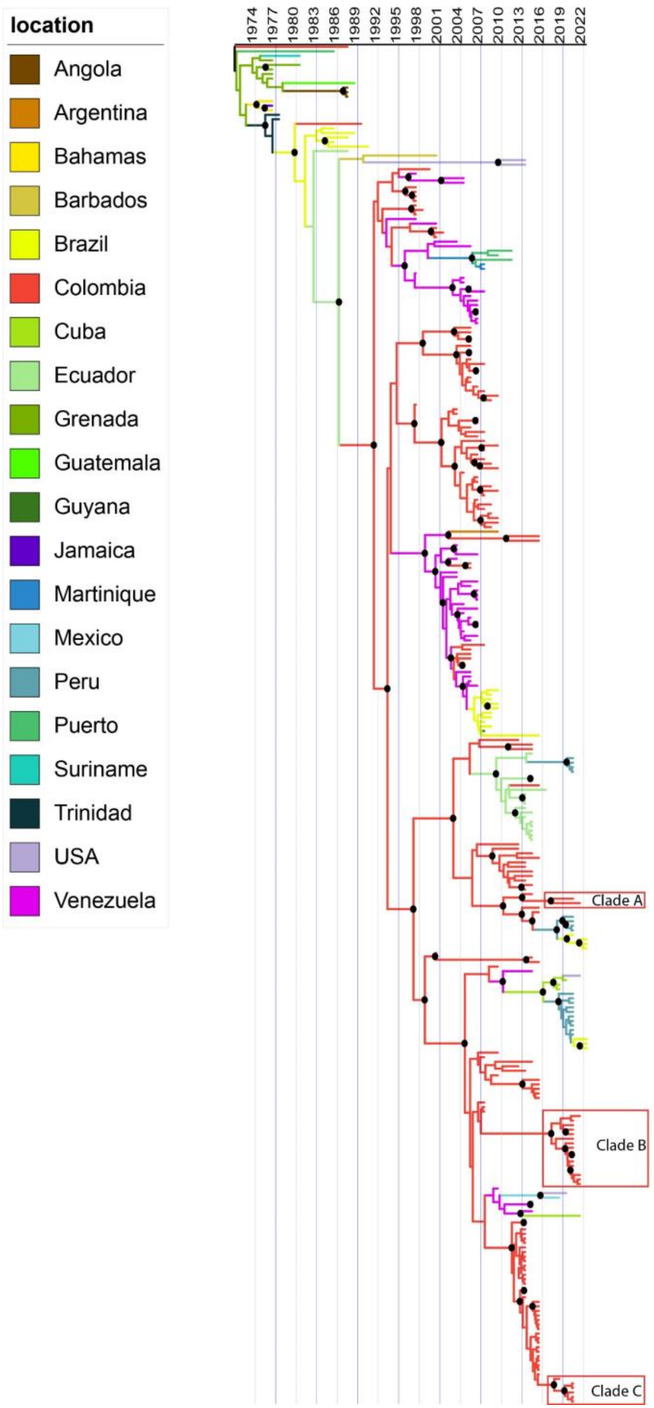


**Figure S7**. Time-scaled Maximum clade credibility (MCC) tree of 300 DENV1 genotype V envelope sequences, including 24 newly-generated sequences. Sequences highlighted in red are from Colombia. Branch colors are based on known or inferred location.

**Table S1**. Primers for DENV1 genotype V multiplex amplicon sequencing. The table lists the DENV-specific primer sequence as well as the sequence of the fusion primers that include the iNEXT universal indexes. The final primer concentrations are also listed.

| Primer Name | Pool | DENV-specific primer (5'-3') | Sequence (with iNEXT fusions primers) 5'->3' | Length | Tm | Final Primer Concentration (µm) |
|---|---|---|---|---|---|---|
| DENV1_1_LEFT | 1 | GAGCAGATCTCTGATGAACAACCA | TCGTCGGCAGCGTCAGATGTGTATAAGAGACAGGAGCAGATCTCTGATGAACAACCA | 57 | 71.5 | 0.56 |
| DENV1_1_RIGHT | 1 | CATGTGTGGCTCTCCCCC | GTCTCGTGGGCTCGGAGATGTGTATAAGAGACAGCATGTGTGGCTCTCCCCC | 52 | 73 | 0.56 |
| DENV1_2_LEFT | 2 | TGACCATGCTCCTCATGCTG | TCGTCGGCAGCGTCAGATGTGTATAAGAGACAGTGACCATGCTCCTCATGCTG | 53 | 72.1 | 0.28 |
| DENV1_2_RIGHT | 2 | CCCAGGTCTCCACTCTTTGTATTT | GTCTCGTGGGCTCGGAGATGTGTATAAGAGACAGCCCAGGTCTCCACTCTTTGTATTT | 58 | 71.3 | 0.28 |
| DENV1_3_LEFT | 1 | ACACGTGGGACTTGGTCTAGAA | TCGTCGGCAGCGTCAGATGTGTATAAGAGACAGACACGTGGGACTTGGTCTAGAA | 55 | 71.7 | 0.56 |
| DENV1_3_RIGHT | 1 | TTGATATTTTAGCTTCAATGCACAGTTTG | GTCTCGTGGGCTCGGAGATGTGTATAAGAGACAGTTGATATTTTAGCTTCAATGCACAGTTTG | 63 | 76.6 | 0.56 |
| DENV1_4_LEFT | 2 | CGTCACCACCATGGCAAAAA | TCGTCGGCAGCGTCAGATGTGTATAAGAGACAGCGTCACCACCATGGCAAAAA | 53 | 72.6 | 0.42 |
| DENV1_4_RIGHT | 2 | GAGCTTGAGGTGTTATGGTTGC | GTCTCGTGGGCTCGGAGATGTGTATAAGAGACAGGAGCTTGAGGTGTTATGGTTGC | 56 | 71.4 | 0.42 |
| DENV1_5_LEFT | 1 | ATAGTCACTGTCCACACTGGGG | TCGTCGGCAGCGTCAGATGTGTATAAGAGACAGATAGTCACTGTCCACACTGGGG | 55 | 71.9 | 0.56 |
| DENV1_5_RIGHT | 1 | CCTGACGTCTGGATTTCTGTCG | GTCTCGTGGGCTCGGAGATGTGTATAAGAGACAGCCTGACGTCTGGATTTCTGTCG | 56 | 72 | 0.56 |
| DENV1_6_LEFT | 2 | ATCTGCTGGTCACATTCAAGAC | TCGTCGGCAGCGTCAGATGTGTATAAGAGACAGATCTGCTGGTCACATTCAAGAC | 55 | 70.8 | 1.13 |
| DENV1_6_RIGHT | 2 | TGTTGACTGGTTTCTCTTTGTCAGT | GTCTCGTGGGCTCGGAGATGTGTATAAGAGACAGTGTTGACTGGTTTCTCTTTGTCAGT | 59 | 70.6 | 1.13 |
| DENV1_7_LEFT | 1 | GCCATGCAAGATCCCCTTCT | TCGTCGGCAGCGTCAGATGTGTATAAGAGACAGGCCATGCAAGATCCCCTTCT | 53 | 72.7 | 0.56 |
| DENV1_7_RIGHT | 1 | CCATGTCAACAGAATCCCTATTCCT | GTCTCGTGGGCTCGGAGATGTGTATAAGAGACAGCCATGTCAACAGAATCCCTATTCCT | 59 | 71.1 | 0.56 |
| DENV1_8_LEFT | 2 | CATCTGTGGGAAAATTGGTACACC | TCGTCGGCAGCGTCAGATGTGTATAAGAGACAGCATCTGTGGGAAAATTGGTACACC | 57 | 71.5 | 0.7 |
| DENV1_8_RIGHT | 2 | TTCAATTCATTTGATATTTGCTTCCACAT | GTCTCGTGGGCTCGGAGATGTGTATAAGAGACAGTTCAATTCATTTGATATTTGCTTCCACAT | 63 | 75.9 | 0.7 |
| DENV1_9_LEFT | 1 | TGTCAGCAGCCATTGGAAAGG | TCGTCGGCAGCGTCAGATGTGTATAAGAGACAGTGTCAGCAGCCATTGGAAAGG | 54 | 72.2 | 0.56 |
| DENV1_9_RIGHT | 1 | GGGTGTAGGAGTCACGCAATTT | GTCTCGTGGGCTCGGAGATGTGTATAAGAGACAGGGGTGTAGGAGTCACGCAATTT | 56 | 71.9 | 0.56 |
| DENV1_10_LEFT | 2 | GACGGCCAAAGAGCATGGAA | TCGTCGGCAGCGTCAGATGTGTATAAGAGACAGGACGGCCAAAGAGCATGGAA | 53 | 72.9 | 0.56 |
| DENV1_10_RIGHT | 2 | TGGTGCCTTCACACAAGTCAAA | GTCTCGTGGGCTCGGAGATGTGTATAAGAGACAGTGGTGCCTTCACACAAGTCAAA | 56 | 71.4 | 0.56 |
| DENV1_11_LEFT | 1 | AGCACAACTACAGACCAGGGTA | TCGTCGGCAGCGTCAGATGTGTATAAGAGACAGAGCACAACTACAGACCAGGGTA | 55 | 71.7 | 0.7 |
| DENV1_11_RIGHT | 1 | AGTCATCAGCATCTTTCTACTCC | GTCTCGTGGGCTCGGAGATGTGTATAAGAGACAGAGTCATCAGCATCTTTCTACTCC | 57 | 69.8 | 0.7 |
| DENV1_12_LEFT | 2 | GGAGAAGTGGACAGTTTTTCATTAGG | TCGTCGGCAGCGTCAGATGTGTATAAGAGACAGGGAGAAGTGGACAGTTTTTCATTAGG | 59 | 70.7 | 0.62 |
| DENV1_12_RIGHT | 2 | GTCAACAATTTTAACATCATGATACCCA | GTCTCGTGGGCTCGGAGATGTGTATAAGAGACAGGTCAACAATTTTAACATCATGATACCCA | 62 | 76.7 | 0.62 |
| DENV1_13_LEFT | 1 | TGGTGGCATCCGTGGAG | TCGTCGGCAGCGTCAGATGTGTATAAGAGACAGTGGTGGCATCCGTGGAG | 50 | 72.7 | 0.62 |
| DENV1_13_RIGHT | 1 | GGAGTGAACTTAGTAGAATGCTGACT | GTCTCGTGGGCTCGGAGATGTGTATAAGAGACAGGGAGTGAACTTAGTAGAATGCTGACT | 60 | 70.5 | 0.62 |
| DENV1_14_LEFT | 2 | CAGAAAACAAAATCTGGGGAAGGA | TCGTCGGCAGCGTCAGATGTGTATAAGAGACAGCAGAAAACAAAATCTGGGGAAGGA | 57 | 71.3 | 0.84 |
| DENV1_14_RIGHT | 2 | TGCCAAAAATACCACACAAAAAGAGT | GTCTCGTGGGCTCGGAGATGTGTATAAGAGACAGTGCCAAAAATACCACACAAAAAGAGT | 60 | 70.2 | 0.84 |
| DENV1_15_LEFT | 1 | GACACACTTACTATACTCCTTAAAGC | TCGTCGGCAGCGTCAGATGTGTATAAGAGACAGGACACACTTACTATACTCCTTAAAGC | 59 | 69.5 | 1.13 |
| DENV1_15_RIGHT | 1 | TCAACAGCTATCACCTGCACTTC | GTCTCGTGGGCTCGGAGATGTGTATAAGAGACAGTCAACAGCTATCACCTGCACTTC | 57 | 71.1 | 1.13 |
| DENV1_16_LEFT | 2 | GGGCCAGTGTCAAAAAGGACTT | TCGTCGGCAGCGTCAGATGTGTATAAGAGACAGGGGCCAGTGTCAAAAAGGACTT | 55 | 72.4 | 0.84 |
| DENV1_16_RIGHT | 2 | TCTTCTTGTTTTTCCGGATCCTGG | GTCTCGTGGGCTCGGAGATGTGTATAAGAGACAGTCTTCTTGTTTTTCCGGATCCTGG | 58 | 70.9 | 0.84 |
| DENV1_17_LEFT | 1 | ATTGCCCAAGCTAAAGCATCACA | TCGTCGGCAGCGTCAGATGTGTATAAGAGACAGATTGCCCAAGCTAAAGCATCACA | 56 | 71.6 | 0.56 |
| DENV1_17_RIGHT | 1 | CTGGCCGCTATGCTGGC | GTCTCGTGGGCTCGGAGATGTGTATAAGAGACAGCTGGCCGCTATGCTGGC | 51 | 73.7 | 0.56 |
| DENV1_18_LEFT | 2 | TTATCCCCAGTGAGAGTTCC | TCGTCGGCAGCGTCAGATGTGTATAAGAGACAGTTATCCCCAGTGAGAGTTCC | 53 | 70.7 | 0.84 |
| DENV1_18_RIGHT | 2 | TGTTGTGACAACATAATCCCAGTC | GTCTCGTGGGCTCGGAGATGTGTATAAGAGACAGTGTTGTGACAACATAATCCCAGTC | 58 | 70.2 | 0.84 |
| DENV1_19_LEFT | 1 | GAAAACGGGTAATCCAATTGAGCA | TCGTCGGCAGCGTCAGATGTGTATAAGAGACAGGAAAACGGGTAATCCAATTGAGCA | 57 | 71.3 | 0.56 |
| DENV1_19_RIGHT | 1 | TCTCTCTCTGGCTCAAAGAGGG | GTCTCGTGGGCTCGGAGATGTGTATAAGAGACAGTCTCTCTCTGGCTCAAAGAGGG | 56 | 71.5 | 0.56 |
| DENV1_20_LEFT | 2 | GGATCACGCTCATTGGACAGAA | TCGTCGGCAGCGTCAGATGTGTATAAGAGACAGGGATCACGCTCATTGGACAGAA | 55 | 72.1 | 0.42 |
| DENV1_20_RIGHT | 2 | GACACTTCTTCTTCCTGCTGCA | GTCTCGTGGGCTCGGAGATGTGTATAAGAGACAGGACACTTCTTCTTCCTGCTGCA | 56 | 71.8 | 0.42 |

| | | | | | | |
|---|---|---|---|---|---|---|
| DENV1_21_LEFT | 1 | CTGGACAAAGGAAGGAGAAAGAAAGA | TCGTCGGCAGCGTCAGATGTGTATAAGAGACAGCTGGACAAAGGAAGGAGAAAGAAAGA | 59 | 71.3 | 0.7 |
| DENV1_21_RIGHT | 1 | CCATCCATAACAGTGCGCTTGA | GTCTCGTGGGCTCGGAGATGTGTATAAGAGACAGCCATCCATAACAGTGCGCTTGA | 56 | 72.2 | 0.7 |
| DENV1_22_LEFT | 2 | GGTGGAGTGACGCTATTCTTCC | TCGTCGGCAGCGTCAGATGTGTATAAGAGACAGGGTGGAGTGACGCTATTCTTCC | 55 | 72.1 | 0.28 |
| DENV1_22_RIGHT | 2 | TGGGAGTGATAACTGTTGTGGC | GTCTCGTGGGCTCGGAGATGTGTATAAGAGACAGTGGGAGTGATAACTGTTGTGGC | 56 | 71.3 | 0.28 |
| DENV1_23_LEFT | 1 | CACCAACATGCTACAATGCTGG | TCGTCGGCAGCGTCAGATGTGTATAAGAGACAGCACCAACATGCTACAATGCTGG | 55 | 72.2 | 0.34 |
| DENV1_23_RIGHT | 1 | TCTATTGCAACAATCCCGTCTACG | GTCTCGTGGGCTCGGAGATGTGTATAAGAGACAGTCTATTGCAACAATCCCGTCTACG | 58 | 70.8 | 0.34 |
| DENV1_24_LEFT | 2 | AAGCAAAGGCCACTAGAGAAGC | TCGTCGGCAGCGTCAGATGTGTATAAGAGACAGAAGCAAAGGCCACTAGAGAAGC | 55 | 71.6 | 0.28 |
| DENV1_24_RIGHT | 2 | TTTCTCTCCCAGTGTTTCCCCT | GTCTCGTGGGCTCGGAGATGTGTATAAGAGACAGTTTCTCTCCCAGTGTTTCCCCT | 56 | 71.6 | 0.28 |
| DENV1_25_LEFT | 1 | GGAGCAGGTCTGGCTTTTTCAT | TCGTCGGCAGCGTCAGATGTGTATAAGAGACAGGGAGCAGGTCTGGCTTTTTCAT | 55 | 72.4 | 0.56 |
| DENV1_25_RIGHT | 1 | CCATAGGTCGCCATTGGGAT | GTCTCGTGGGCTCGGAGATGTGTATAAGAGACAGCCATAGGTCGCCATTGGGAT | 54 | 72.3 | 0.56 |
| DENV1_26_LEFT | 2 | TGCTGGGCTGAAGAAAGTTACT | TCGTCGGCAGCGTCAGATGTGTATAAGAGACAGTGCTGGGCTGAAGAAAGTTACT | 55 | 71.4 | 0.48 |
| DENV1_26_RIGHT | 2 | TTTCCTGTTCCACATGAAACCCA | GTCTCGTGGGCTCGGAGATGTGTATAAGAGACAGTTTCCTGTTCCACATGAAACCCA | 57 | 71.1 | 0.48 |
| DENV1_27_LEFT | 1 | AAACACGGAGGGATGCTAGTG | TCGTCGGCAGCGTCAGATGTGTATAAGAGACAGAAACACGGAGGGATGCTAGTG | 54 | 71.6 | 0.56 |
| DENV1_27_RIGHT | 1 | ACATCCCATGGTTTTGTGAGCA | GTCTCGTGGGCTCGGAGATGTGTATAAGAGACAGACATCCCATGGTTTTGTGAGCA | 56 | 71.3 | 0.56 |
| DENV1_28_LEFT | 2 | CATGGATCATATGAGGTCAAGCCA | TCGTCGGCAGCGTCAGATGTGTATAAGAGACAGCATGGATCATATGAGGTCAAGCCA | 57 | 71.7 | 0.28 |
| DENV1_28_RIGHT | 2 | TCTCTGTGCACGAGATCCCA | GTCTCGTGGGCTCGGAGATGTGTATAAGAGACAGTCTCTGTGCACGAGATCCCA | 54 | 71.9 | 0.28 |
| DENV1_29_LEFT | 1 | AGGAGCAGTGTTCGTTGATGAAA | TCGTCGGCAGCGTCAGATGTGTATAAGAGACAGAGGAGCAGTGTTCGTTGATGAAA | 56 | 71.3 | 0.56 |
| DENV1_29_RIGHT | 1 | TTATTCTTGTGTCCCATCCGGC | GTCTCGTGGGCTCGGAGATGTGTATAAGAGACAGTTATTCTTGTGTCCCATCCGGC | 56 | 71.3 | 0.56 |
| DENV1_30_LEFT | 2 | GGACTGCACAAACTTGGATACA | TCGTCGGCAGCGTCAGATGTGTATAAGAGACAGGGACTGCACAAACTTGGATACA | 55 | 71.4 | 0.7 |
| DENV1_30_RIGHT | 2 | CCAACCAGTCAAGAACTCTCTCAG | GTCTCGTGGGCTCGGAGATGTGTATAAGAGACAGCCAACCAGTCAAGAACTCTCTCAG | 58 | 71.4 | 0.7 |
| DENV1_31_LEFT | 1 | GGAGGTCCAACTAATAAGACAAATGGA | TCGTCGGCAGCGTCAGATGTGTATAAGAGACAGGGAGGTCCAACTAATAAGACAAATGGA | 60 | 70.8 | 0.56 |
| DENV1_31_RIGHT | 1 | AGTTTCTCTCAGGCTCCATCC | GTCTCGTGGGCTCGGAGATGTGTATAAGAGACAGAGTTTCTCTCAGGCTCCATCC | 55 | 71 | 0.56 |
| DENV1_32_LEFT | 2 | AGGGAAATAGTGGTGCCATGC | TCGTCGGCAGCGTCAGATGTGTATAAGAGACAGAGGGAAATAGTGGTGCCATGC | 54 | 71.9 | 0.42 |
| DENV1_32_RIGHT | 2 | TGTCAAGCCTATCAGGGATCCA | GTCTCGTGGGCTCGGAGATGTGTATAAGAGACAGTGTCAAGCCTATCAGGGATCCA | 56 | 71.4 | 0.42 |
| DENV1_33_LEFT | 1 | GGATGGAGGACAAAACTCATGTATCC | TCGTCGGCAGCGTCAGATGTGTATAAGAGACAGGGATGGAGGACAAAACTCATGTATCC | 59 | 71.3 | 0.56 |
| DENV1_33_RIGHT | 1 | CTTGCTCAATCCGTGGCTTTC | GTCTCGTGGGCTCGGAGATGTGTATAAGAGACAGCTTGCTCAATCCGTGGCTTTC | 55 | 72 | 0.56 |
| DENV1_34_LEFT | 2 | CGAAGGAGCACTCTGGTAAGTC | TCGTCGGCAGCGTCAGATGTGTATAAGAGACAGCGAAGGAGCACTCTGGTAAGTC | 55 | 72.2 | 0.65 |
| DENV1_34_RIGHT | 2 | TGGTCTCTCCCAGCGTCAATAT | GTCTCGTGGGCTCGGAGATGTGTATAAGAGACAGTGGTCTCTCCCAGCGTCAATAT | 56 | 71.5 | 0.65 |

**Table S2. Clinical and laboratory-confirmed DENV cases in two Colombian departments: Valle del Cauca and Risaralda**. Serum from patients with acute febrile symptoms (Risaralda) and clinically diagnosed DENV (Valle del Cauca) were analyzed using the NS1/IgM/IgG rapid test and RT-qPCR. Samples that were positive for DENV RNA underwent typing RT-qPCR, and DENV1 samples underwent full genome sequencing.

| Department | Samples | NS1/IgM rapid test | RT-qPCR | Serotype | | | | Sequenced |
|---|---|---|---|---|---|---|---|---|
| | | | | DENV1 | DENV2 | DENV3 | DENV4 | |
| Valle del Cauca | 201 | 96 | 79 | 42 | 20 | 4 | 0 | 17 |
| Risaralda | 178 | 20 | 20 | 7 | 1 | 0 | 0 | 7 |

**Table S3**. Twenty-four new full-length DENV1 genotype V sequences were generated from two Colombian departments from 2019-2022.

| GenBank Accession | Department | Collection Date | Sample ID | Coverage |
|---|---|---|---|---|
| PP957577 | Valle del Cauca | 8/15/2021 | VAC_DFC059_2021 | 100% |
| PP957578 | Valle del Cauca | 8/15/2021 | VAC_DFC063_2021 | 100% |
| PP957579 | Valle del Cauca | 8/24/2021 | VAC_DFC087_2021 | 100% |
| PP957580 | Valle del Cauca | 8/28/2021 | VAC_DFC088_2021 | 100% |
| PP957581 | Valle del Cauca | 8/30/2021 | VAC_DFC096_2021 | 100% |
| PP957582 | Valle del Cauca | 9/6/2021 | VAC_DFC112_2021 | 100% |
| PP957583 | Valle del Cauca | 9/6/2021 | VAC_DFC114_2021 | 100% |
| PP957584 | Valle del Cauca | 10/16/2021 | VAC_DFC161_2021 | 100% |
| PP957585 | Valle del Cauca | 10/17/2021 | VAC_DFC162_2021 | 100% |
| PP957586 | Valle del Cauca | 10/29/2021 | VAC_DFC177_2021 | 100% |
| PP957591 | Valle del Cauca | 5/11/2022 | VAC_DFC263_2022 | 100% |
| PP957592 | Valle del Cauca | 11/11/2022 | VAC_DFC264_2022 | 100% |
| PP957593 | Valle del Cauca | 11/12/2022 | VAC_DFC265_2022 | 99% |
| PP957590 | Valle del Cauca | 7/24/2022 | VAC_DFC245_2022 | 100% |
| PP957589 | Valle del Cauca | 12/12/2021 | VAC_DFC238_2021 | 100% |
| PP957588 | Valle del Cauca | 12/3/2021 | VAC_DFC222_2021 | 100% |
| PP957587 | Valle del Cauca | 11/30/2021 | VAC_DFC217_2021 | 100% |
| PP957573 | Risaralda | 3/3/2021 | RIS_AFI 031_2021 | 100% |
| PP957574 | Risaralda | 6/28/2021 | RIS_AFI 082_2021 | 95% |
| PP957575 | Risaralda | 8/5/2021 | RIS_AFI 101_2021 | 96% |
| PP957576 | Risaralda | 4/21/2022 | RIS_AFI 224_2022 | 100% |
| OM654348 | Risaralda | 1/25/2021 | RIS_OM654348_2021 | 99.70% |
| MZ773407 | Risaralda | 9/21/2019 | RIS_MZ773407_2019 | 100% |
| OM654347 | Risaralda | 1/25/2021 | RIS_OM654347_2021 | 100% |

**Table S4.** Nested sampling results for model selection for CDS dataset and Protein E Data set

| CDS Data set | | | | |
|---|---|---|---|---|
| **Model** | **Marginal Likelihood** | **SD** | **Information** | **ML Difference** |
| Strict_Constant | -43656.55606 | sqrt(H/N)=(2.0)=?=SD=(1.9) | 267.6 | 310.8 |
| Strict _Exponential | -43792.87992 | sqrt(H/N)=(2.0)=?=SD=(2.0) | 269.1 | 447.1 |
| Strict _BS | -43662.96482 | sqrt(H/N)=(1.9)=?=SD=(1.9) | 268.8 | 317.2 |
| Strict _EBS | -43658.8195 | sqrt(H/N)=(2.0)=?=SD=(2.0) | 272.8 | 313.0 |
| Relax Lognormal_Constant | -43361.37548 | sqrt(H/N)=(2.0)=?=SD=(2.0) | 284.8 | 15.6 |
| Relax Lognormal_Exponential | -43377.10127 | sqrt(H/N)=(2.0)=?=SD=(2.0) | 284.8 | 31.3 |
| Relax Lognormal_BS* | -43345.78699 | sqrt(H/N)=(2.0)=?=SD=(2.0) | 284.5 | |
| Relax Lognormal_EBS | -43359.01884 | sqrt(H/N)=(2.0)=?=SD=(1.9) | 284.6 | 13.2 |
| **Protein E Data set** | | | | |
| **Model** | **Marginal Likelihood** | **SD** | **Information** | **ML Difference** |
| Strict_Constant | -8516.409458 | sqrt(H/N)=(8.4)=?=SD=(8.3) | 1119.2 | 46.7 |
| Strict _Exponential | -8530.255461 | sqrt(H/N)=(8.3)=?=SD=(8.5) | 1104.8 | 60.5 |
| Strict _BS* | -8469.750158 | sqrt(H/N)=(8.2)=?=SD=(8.2) | 1097.7 | |
| Strict _EBS | -8515.660103 | sqrt(H/N)=(8.4)=?=SD=(8.6) | 1141.3 | 45.9 |
| Relax Lognormal_Constant | -8517.495131 | sqrt(H/N)=(8.5)=?=SD=(8.6) | 1146.1 | 47.7 |
| Relax Lognormal_Exponential | -8509.69687 | sqrt(H/N)=(8.6)=?=SD=(9.1) | 1192.2 | 39.9 |
| Relax Lognormal_BS | -8496.329628 | sqrt(H/N)=(8.6)=?=SD=(8.4) | 1193.8 | 26.6 |
| Relax Lognormal_EBS | -8737.887388 | sqrt(H/N)=(8.8)=?=SD=(9.0) | 1228.2 | 268.1 |

*Model with the highest likelihood value. Minimum expected difference for whole genomes dataset: 5.7 and Minimum expected difference for Full-length E protein data set: 23.6.

**Table S5**. **Summary of the Time to the Most Recent Common Ancestor (TMRCA) for nodes of interest.** The CDS data was analyzed with a best fit model using a relaxed molecular clock, and the envelope gene dataset was analyzed with a best fit model using a strict molecular clock. NC=not calculated, since the CDS data set does not include the sequences from Santander used to identify clade B*.

| **Data set** | **Clade** | **MRCA date** | **HPDI** | |
|---|---|---|---|---|
| **CDS** | Clade A | 2017.88 | 2015.73 | 2019.84 |
| | Clade A* | 2014.21 | 2012.78 | 2015.02 |
| | Clade B | 2016.74 | 2015.01 | 2018.38 |
| | Clade B* | NC | NC | NC |
| | Clade C | 2018.78 | 2017.95 | 2019.21 |
| | Clade C* | 2013.28 | 2012.51 | 2013.80 |
| ***ENV*** | Clade A | 2018.07 | 2016.17 | 2019.89 |
| | Clade A* | 2013.86 | 2012.12 | 2015.10 |
| | Clade B | 2018.11 | 2016.40 | 2019.50 |
| | Clade B* | 2007.78 | 2007.02 | 2008.23 |
| | Clade C | 2018.52 | 2017.18 | 2019.25 |
| | Clade C* | 2016.10 | 2015.58 | 2016.25 |
